# CCL20–CCR6 Signaling as a Prognostic Biomarker and Therapeutic Target in Temozolomide-Resistant Glioblastoma

**DOI:** 10.64898/2026.08.26.746721

**Authors:** Ryan Green, Karthick Mayilsamy, Erin Anglin, Kristina Tosi, Sashank Bikkasani, Eleni Markoutsa, Parthvi Bharatkumar Patel, Tiara Wolf, Jennifer Guergues, Stanley M. Stevens, Ganesh Halade, Shyam Mohapatra, Subhra Mohapatra

## Abstract

Glioblastoma remains highly lethal, with median survival of ∼15 months. Resistance to temozolomide is ubiquitous, yet its mechanisms are incompletely understood. Here, we identify the CCL20-CCR6 chemokine axis as a stress-responsive survival pathway limiting therapeutic efficacy. Targeting CCL20–CCR6 in combination with temozolomide and cannabidiol was evaluated using clinical datasets, GBM cell lines, tumor organoids, and a syngeneic CT-2A mouse model integrating proteomic and lipidomic profiling. Low CCL20 expression was associated with improved survival, supporting its prognostic relevance. Across models, TMZ alone or with CBD induced CCL20 expression while exerting limited antitumor activity. Targeted disruption of CCL20-CCR6 signaling using dendrimer-delivered shRNA enhanced therapeutic response in murine models and GBM organoids. Multi-omic analyses revealed that CCL20 inhibition reprograms the tumor microenvironment and induces mitochondrial dysfunction, resulting in elevated reactive oxygen species (ROS) and tumor cell death. This effect was accompanied by accumulation of 17-hydroxydocosahexaenoic acid and activation of oxidative stress-associated cytotoxic pathways. Functional assays confirmed that CCL20 blockade selectively amplifies mitochondrial ROS beyond levels induced by TMZ alone potentiating TMZ efficacy by promoting mitochondrial oxidative stress. Targeting this axis represents a promising strategy to overcome chemoresistance and positions CCL20 as both a prognostic biomarker and a therapeutic vulnerability in GBM.

**Key Points:**

1. CCL20-CCR6 is a clinically relevant adaptive resistance pathway in GBM
2. Targeting CCL20-CCR6 significantly enhances TMZ-based therapy
3. Therapeutic benefit is driven by mitochondrial dysfunction and oxidation stress

**Importance of this study:** Our study identifies a previously underappreciated role for the chemokine CCL20 in regulating metabolism and mitochondrial function in GBM. In addition, we show that targeting the CCL20/CCR6 axis can significantly influence therapeutic outcomes. Prior studies have reported elevated CCL20 expression in glioblastoma relative to normal tissue, as well as its induction following TMZ treatment. Building on these observations, our findings demonstrate that inhibition of CCL20 may act synergistically with TMZ by disrupting redox homeostasis and promoting mitochondrial dysfunction.

These results have important translational implications. Validating CCL20 as a therapeutic target to enhance TMZ efficacy supports further development of small-molecule inhibitors of the CCL20/CCR6 pathway. Moreover, investigating oxidative stress mechanisms downstream of CCL20 may uncover additional strategies to circumvent CCR6-dependent survival pathways, ultimately improving responses to TMZ and advancing GBM treatment.

## Introduction

Gliomas are the most common malignant primary brain tumors, with an annual incidence of ∼6 per 100,000 individuals in the United States^1,2^. These tumors comprise a heterogenous group, including astrocytoma, ependymoma, oligodendroglioma, and glioblastoma (GBM). GBM (WHO grade-4, IDH wild-type) accounts for ∼45-50% of cases and is the most aggressive subtype^3,4^. Despite advances in surgery, radiation therapy, and immunotherapies (anti-CTLA-4, PD-1, CAR-T), patient outcomes remain poor, largely due to an immunosuppressive tumor microenvironment (TME) and limited immune infiltration. Median survival is 9-18 months, with a 5-year survival of 5-10%^5–7^. The standard of care (Stupp protocol) combines surgical resection with radiotherapy and concurrent/adjuvant Temozolomide (TMZ)^8,9^, improving 2-yr survival compared to radiation alone^9^. Subsequent meta-analyses have confirmed a survival benefit (HR 0.63)^10^. Nevertheless, long-term outcomes remain dismal, largely due to intrinsic and acquired TMZ resistance.

TMZ induces DNA damage primarily through DNA alkylation (O^6^-methylguanine), leading to DNA mismatch, strand breaks, and cytotoxicity^11^. The endogenous DNA repair enzyme methylguanine methyltransferase (MGMT) can reverse this lesion^12^. MGMT promoter methylation, resulting in gene silencing, is a key determinant of TMZ sensitivity^13^. However, many GBMs lacking MGMT promoter methylation exhibit resistance, reflecting additional mechanisms of therapeutic evasion. These include activation of pro-growth signaling pathways (e.g., PI3K, Wnt, EGFR, Ras-ERK, MAPK), upregulation of antioxidant systems such as superoxide dismutase 2, enhanced DNA repair through base excision repair or mismatch repair pathways, glioma stem cell maintenance, autophagy, and evasion of apoptosis ^14–16^. Mitochondria also play a crucial role in TMZ resistance through mechanisms including modulation of reactive oxygen species (ROS), metabolic reprogramming, and apoptosis regulation^17^. Together, these mechanisms underscore the need for therapeutic strategies that sensitize tumors to TMZ.

The chemokine CCL20 and its receptor CCR6 regulate immune cell recruitment, inflammation, and tumor progression^18–20^. In the brain, CCL20 has been implicated in neuroinflammatory response following traumatic brain injury^21^. Emerging evidence suggests that CCL20 expression correlates with poor clinical outcomes in GBM^22–24^ and activates pathways overlapping with TMZ resistance, including ERK, PI3K, and NFκB signaling. In GBM, CCL20 promotes tumor progression by remodeling the TME, recruiting immunosuppressive myeloid cells, and supporting glioma stem cell maintenance^25,26^. Although the interaction between TMZ and CCL20 remains incompletely understood, *in vivo* studies demonstrate increased CCL20 expression in GBM following TMZ treatment^23^. Notably, TMZ-induced stress responses activate p38 MAPK and NF-kB^27–30^ signaling pathways, which are known regulators of CCL20 expression. Collectively, these reports led us to postulate that CCL20 is a key driver of TMZ resistance and that targeting the CCL20-CCR6 axis may enhance therapeutic response.

Cannabidiol (CBD), a non-psychoactive cannabinoid, has emerged as a potential adjunct therapy with anti-tumor activity across multiple cancers. CBD interacts with a broad range of molecular targets^31,32^ and modulates proliferation, apoptosis, angiogenesis, and TME^33–36^. Preclinical studies suggest synergy between TMZ and CBD in glioma, although clinical evidence remains limited. A recent phase 1b placebo-controlled trial evaluating cannabinoids (CBD+ Δ^9^-tetrahydrocannabinol, THC) with TMZ in recurrent GBM demonstrates safety and potential survival benefit (n=20)^37^, but larger studies are needed^38–40^, and the distinct contributions of CBD versus THC remain unclear^41,42^. Based on our preliminary observations that treatment of tumor cells with TMZ and/or CBD enhances CCL20 expression, we hypothesized that CCL20 signaling promotes TMZ resistance and that its inhibition would enhance therapeutic response. To test this, we evaluated the CCL20-CCR6 axis using clinical datasets and tumor tissue microarrays, *in vitro* GBM models, organoid cultures, and *in vivo* mouse models. Integrated proteomic and lipidomic analyses were performed to define underlying mechanisms. Our findings identify CCL20 as a key driver of TMZ resistance through modulation of immune and mitochondrial pathways and support targeting the CCL20-CCR6 axis as a strategy to improve GBM therapy.

## Results

### CCL20 is upregulated in glioma and associated with poor clinical outcomes

To investigate the role of CCL20 in glioma, we first assessed its expression in patient-derived tumor tissues. Tissue microarrays containing GBM specimens alongside normal cerebrum controls were immunostained for CCL20 (**Fig 1A**). Quantitative analysis across tumor cases and controls revealed significantly elevated CCL20 protein levels in GBM tissues compared with normal controls. To validate these findings in a larger cohort, we compared CCL20 mRNA expression in GBM tumors within the TCGA database, with normal brain tissue from the Genotype-Tissue Expression (GTEx) database, via the Gene Expression Profiling Interactive Analysis (GEPIA2) platform (**Fig 1B**)^43^. This analysis demonstrated more than a twofold increase in CCL20 expression in GBM, confirming transcriptional upregulation. Although higher mean CCL20 expression was also observed in low grade gliomas, the increase relative to normal tissue was not statistically significant (**Fig S1**). These findings suggest that CCL20 expression may correlate with tumor grade. We next assessed the clinical relevance of CCL20 expression by performing survival analyses using datasets from the NIH Genomic Data Commons (GDC) (**Fig 1C, S2**). Across all glioma patients, stratification by CCL20 expression (cutoff of 0.5 uqFPKM) revealed significantly improved overall survival in the low-expression group. Given that TMZ is the most commonly administered therapy for glioma, we conducted a subgroup analysis of TMZ-treated patients. This analysis revealed a dose-dependent inverse relationship between CCL20 expression and survival: patients with undetectable CCL20 levels exhibited the longest survival, whereas those with very high CCL20 expression (>20 uqFPKM) had the shortest survival. In contrast, CCL20 expression did not significantly correlate with survival in patients treated with Bevacizumab, the second most frequently used therapeutic agent, suggesting a treatment-specific association. Finally, we evaluated whether CCL20 is associated with tumor recurrence. Differential gene expression analysis of RNA-sequencing data from matched primary and recurrent GBM samples (NIH GDC) revealed a greater than six-fold increase in CCL20 expression in recurrent tumors (**Fig1D**).

**Figure 1.**
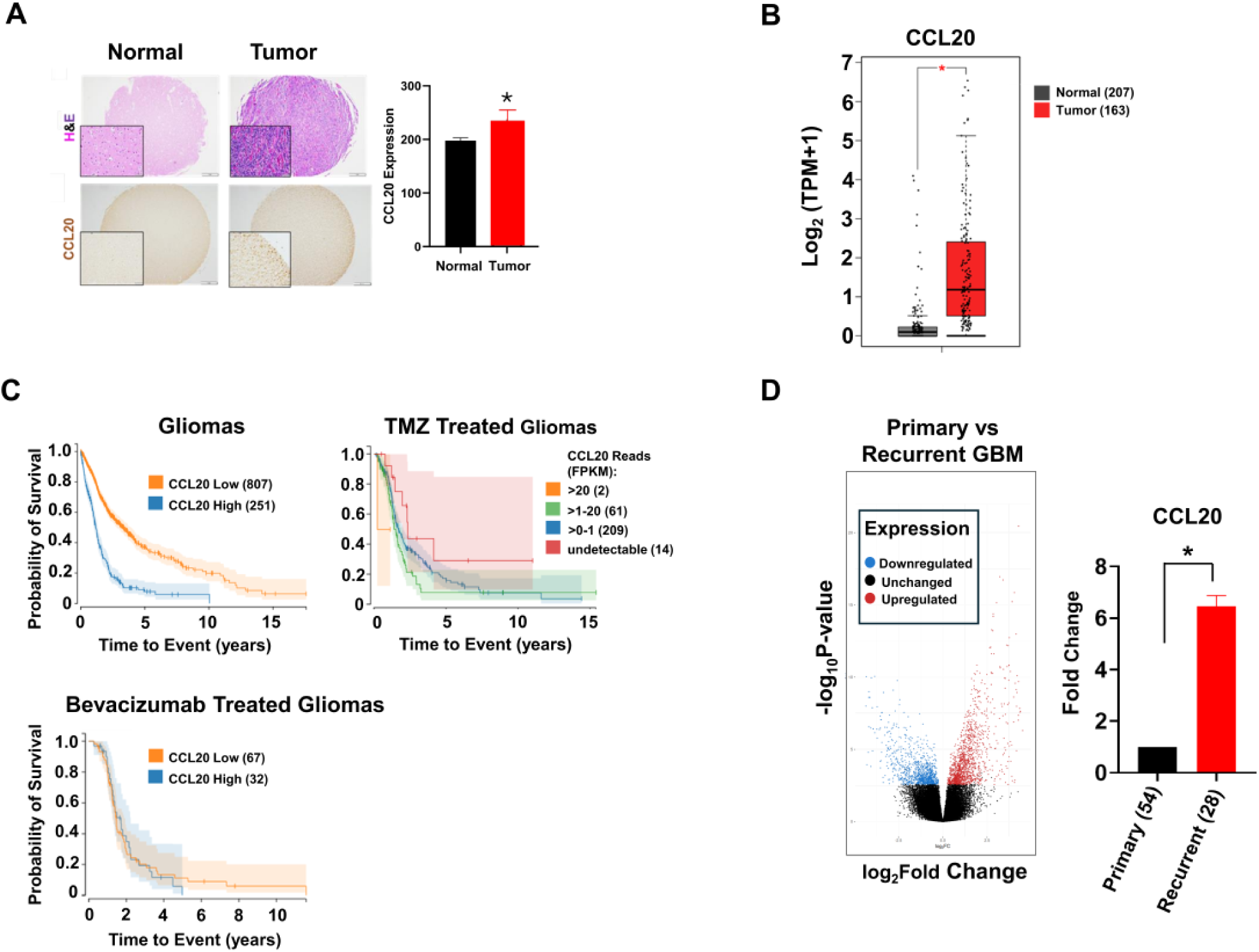
CCL20 is associated with unfavorable outcomes in glioma. CCL20 expression was evaluated by immunostaining and RNA sequencing in gliomas vs normal brain tissue, and by RNA sequencing in recurrent vs primary gliomas. **(A)** Human control brain vs GBM tissue sections were immunostained for CCL20. The intensity of 3,3′-Diaminobenzidine (DAB) signal was quantified using ImageJ (Student’s t test, n=5 controls and 12 GBM patients. *=p≤0.05). **(B)** Gene expression analysis of human brain tissue (normal vs tumor) was performed using the GEPIA2. CCL20 expression was compared between normal brain samples (n=207) and GBM samples (n=163) using ANOVA with Benjamini - Hochberg false discovery rate correction (*=p≤0.01). **(C)** The NIH Genomic Data Commons was queried for survival data in glioma patients stratified by CCL20 expression: high (≥0.5 uqFPKM) vs low (< 0.5 uqFPKM). Survival differences were assessed using the log-rank test (p≤0.05). **(D)** Differential gene expression analysis was performed on RNA sequencing datasets from NIH dbGAP, comparing recurrent (n=28) and primary (n=54) glioma tumor samples.

### TMZ and CBD Combination treatment synergistically enhances cytotoxicity and upregulates CCL20 signaling in glioblastoma cells

To investigate the role of CCL20 in treatment response, we utilized a panel of murine and human GBM cell lines (CT-2A, GL261, SMA560, U87, U251) representing diverse oncogenic driver mutations (**S Table 1)**. Given the well-established resistance of these models to TMZ (IC50 range: 100µM-2mM) and the limited clinical efficacy of TMZ monotherapy, we first assessed their sensitivity to TMZ and CBD. Viability assays confirmed limited responsiveness to TMZ across all cell lines, with IC50 values of ≥1mM (**Fig 2A-B**). In contrast, sensitivity to CBD varied, with IC50 values ranging from 35µM in CT-2A to >100µM in U87 cells. Notably, co-treatment with a suboptimal dose of TMZ (500µM) enhanced the cytotoxic effects of CBD. To quantitatively evaluate drug interactions, combination indices were calculated and demonstrated synergistic effects of TMZ+CBD in all cell lines except U87, where the interaction was additive (**Fig 2B**). These results suggest that CBD can sensitize GBM cells to TMZ despite intrinsic resistance. Given the association of CCL20 with tumor recurrence and patient survival, we next examined whether treatment modulates CCL20 expression. Quantitative real-time PCR analysis revealed upregulation of CCL20 following TMZ+CBD treatment in CT-2A, GL261, and U251 cells. In U87 cells, TMZ monotherapy alone was sufficient to induce CCL20 expression (**Fig 2C**). To further investigate the role of CCL20 in TMZ response, CCL20 knockout (KO) cell lines were generated in CT-2A and GL261 cells using CRISPR/Cas9 genome editing (**Fig S3A**). TMZ sensitivity was then compared between parental and KO cells (**Fig 2D**). Loss of CCL20 increased sensitivity to TMZ in both models suggesting that CCL20 contributes to TMZ resistance. Furthermore, CCL20 KO CT-2A exhibited a slower growth rate than WT CT-2A *in vivo* when implanted subcutaneously into C57 BL/6 mice (**Fig S3B**).

**Figure 2.**
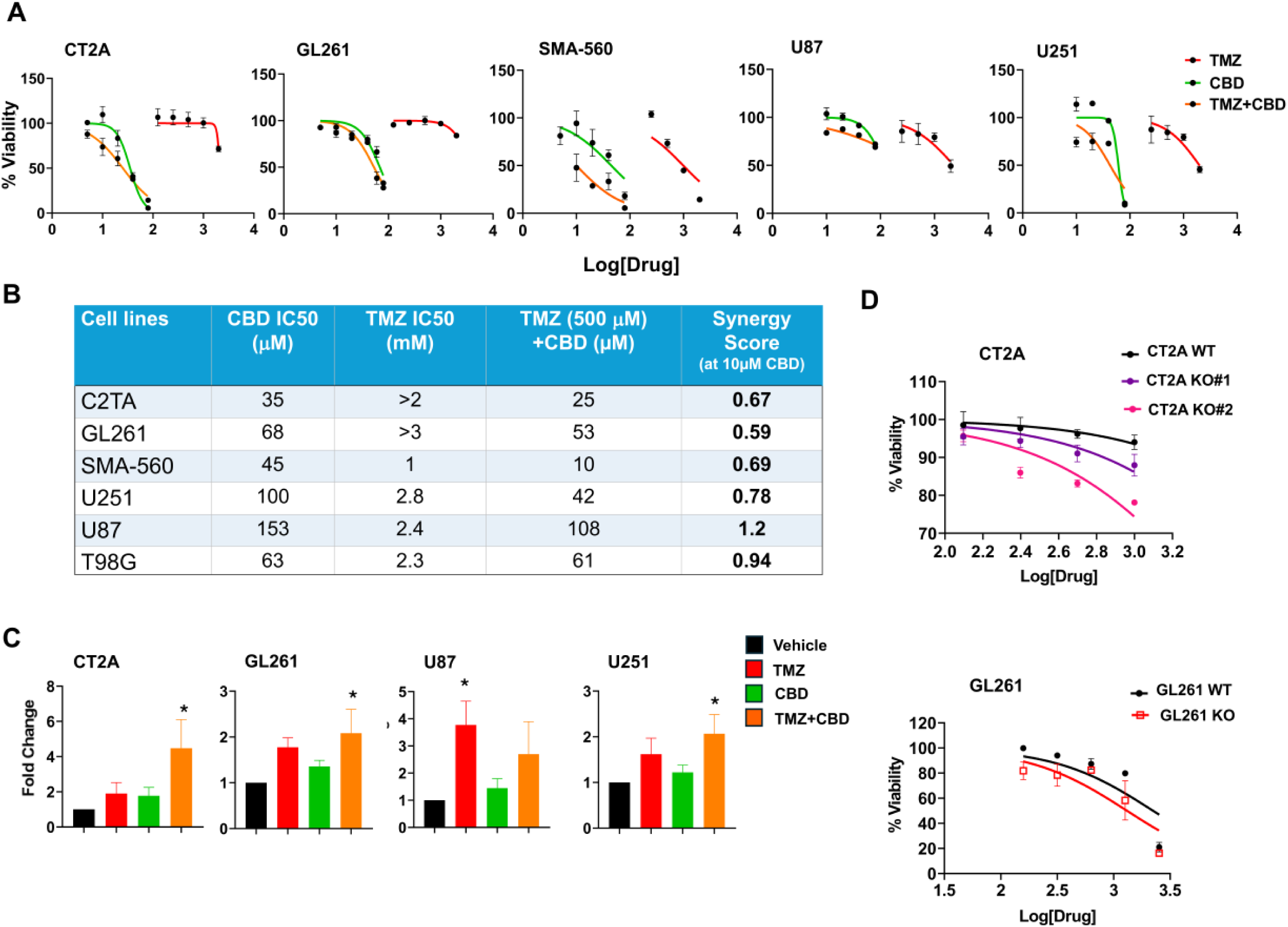
CBD sensitizes GBM to TMZ while inducing CCL20 expression. Effects of TMZ and CBD, alone or in combination, were evaluated in GBM cell lines. **(A)** Dose-response curves of murine and human cell lines treated with TMZ, CBD, or their combination. **(B)** Calculated IC50 values and drug synergy (combination index) derived from the dose-response data shown in (A) (GraphPad Prism, CompuSyn analyses). **(C)** Relative CCL20 expression following drug treatment, measured by qPCR (n=3; one-way ANOVA with Dunnet post hoc test *=p≤0.05). **(D)** TMZ Dose-response curves comparing WT CT2A and GL261 cell lines with CRISPR/Cas9-mediated CCL20 knockout lines.

### CCL20–CCR6 blockade potentiates combination therapy and reduces tumor progression in GBM

Given the established association between elevated CCL20 levels and poor clinical outcomes, we hypothesized that targeting the CCL20–CCR6 signaling axis may enhance sensitivity to TMZ and CBD. To test this hypothesis in vivo, we employed a syngeneic murine GBM model generated by subcutaneous inoculation of CT-2A cells into the hind flanks of C57BL/6 mice. To selectively disrupt CCL20/CCR6 signaling, we developed a G4-PAMAM) dendrimer-based nanosystem (DPX) complexed with shRNA plasmids targeting both CCL20 and CCR6. This platform enables efficient nucleic acid delivery via electrostatic interactions and has demonstrated translational potential for brain tumors via “nose-to-brain” administration, as previously described^44^. PAMAM dendrimers offer several advantages, including high cargo-loading capacity, multiple functionalization sites, and favorable safety profiles in phase I and II clinical trials.

Once tumors reached approximately 6 mm in diameter, mice were randomized to receive vehicle, CBD (25mg/kg oral), TMZ (20mg/kg IP), or combination therapy every other day, with or without the addition of the CCL20/CCR6-targeting DPX (shDPX) (**Fig 3A**). The shDPX formulation (10 ug of plasmid encoding shCCL20 and shCCR6) or scrambled control was administered IP twice weekly. Vehicle-treated mice exhibited rapid tumor progression, whereas the triple combination therapy group (TMZ+CBD+shDPX) showed the most pronounced inhibition of tumor growth. Notably, the addition of ShDPX significantly enhanced the efficacy of combined CBD and TMZ treatment, while no significant benefit was observed when shDPX was combined with either agent alone (**Fig 3B**). At study endpoint (day 30), tumors were harvested for molecular and histological analyses. Tumor weights were lowest in the triple combination group compared with all other treatments (**Fig S4A**).

**Figure 3.**
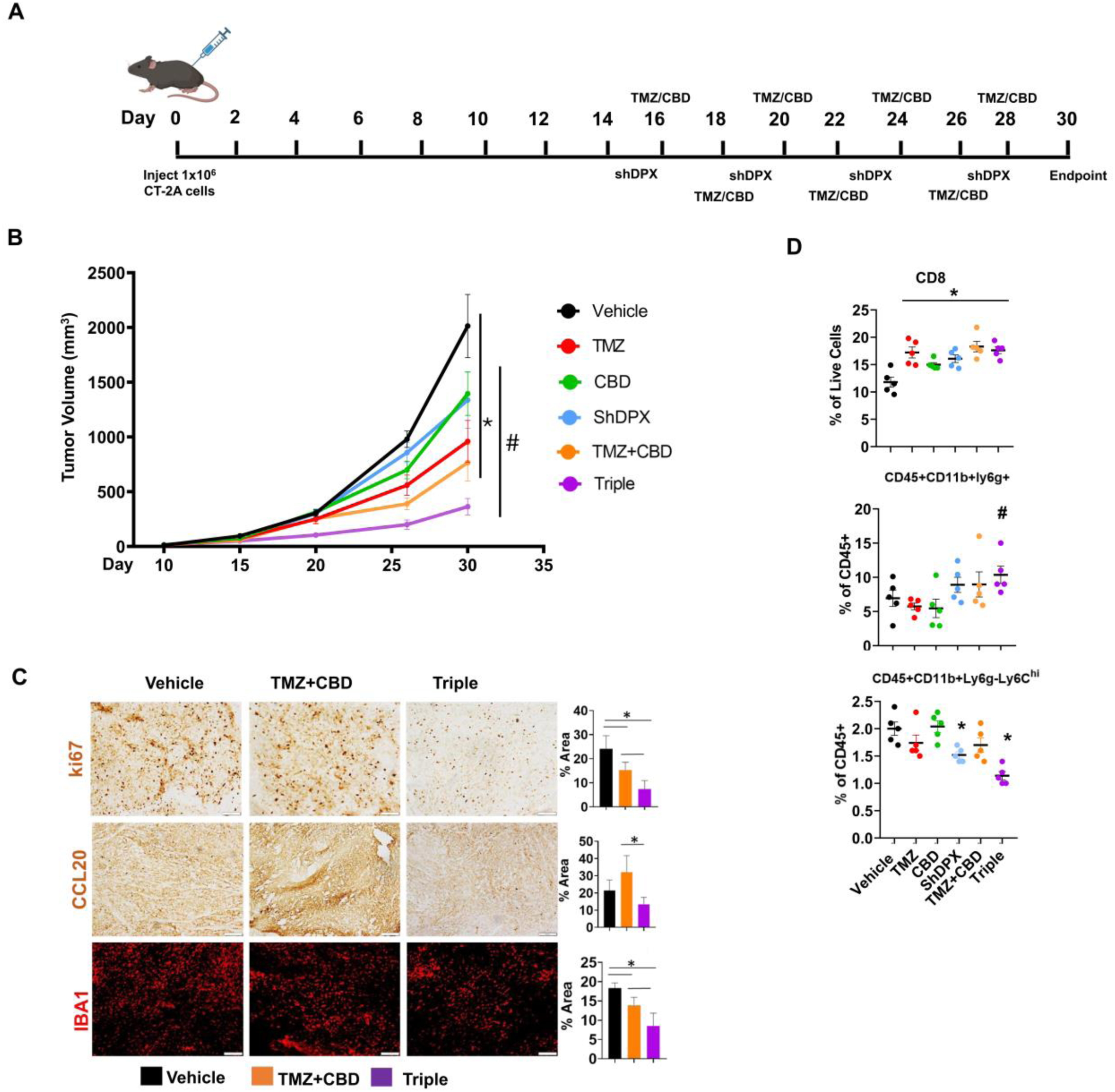
Inhibition of CCL20/CCR6 enhances the efficacy of TMZ + CBD combination therapy. The effectiveness of single agents and combination treatments were evaluated in vivo using a subcutaneous CT-2A flank tumor model. **(A)** Experimental timeline depicting the treatment schedule. **(B)** Growth of subcutaneous CT-2A tumors in mice treated with single agents or combination therapies. C57BL/6 mice (n=5/group) were inoculated subcutaneously in the flank with 1×10^6^ CT2A cells on day 0. Treatments were initiated on day 15 and administered every third day. Treatment groups included vehicle, CBD (25mg/kg, oral gavage), TMZ (20mg/kg, IP), or combination therapies with or without shDPX (IP). Statistical significance: * vs triple, # vs vehicle (one-way ANOVA with Holm Sidack p≤0.05). **(C)** Tumors were harvested, fixed and cryosectioned for immunostaining of Ki67, IBA1, and CCL20. Signal intensity was quantified using ImageJ (Student’s t test, n=3 *=p≤0.05). **(D)** Immunophenotyping of splenocytes was performed by flowcytometry (n=5/group). Statistical comparisons: * vs vehicle, #vs TMZ, p<0.05.

Immunohistochemical (IHC) staining of cryosectioned tumors was performed for CCL20, the proliferation marker Ki67, and the glial activation marker ionized calcium-binding adapter molecule 1 (IBA1) (**Fig3C**). Treatment with TMZ+CBD significantly reduced Ki67 expression, with a further decrease observed in the triple combination group, indicating enhanced suppression of tumor proliferation. In contrast, CCL20 expression was elevated in tumors from TMZ+CBD-treated mice but returned to baseline in the triple combination group. IBA1 staining revealed a substantial reduction in tumor-associated macrophage infiltration in both the TMZ+CBD and triple combination groups, with the most pronounced reduction observed in the latter (**Fig 3C**). Spleens were also collected for immunophenotyping by flow cytometry (**Fig S4B-C**). All treatment groups showed a significant increase in CD3+CD8+ T cells (**Fig 3D**).

Notably, both the shDPX combination and triple combination treatments significantly decreased the frequency of CD11b^+^Ly6^-^Ly6C^hi^ monocytic myeloid-derived suppressor cells (m-MDSCs), while the triple combination additionally increased CD11b^+^Ly6G^+^ granulocytic MDSCs (g-MDSCs). These shifts suggest that a reduction in m-MDSCs, together with an increase in g-MDSCs, may contribute to enhanced anti-tumor responses through modulation of the TME. Collectively, these findings suggest that targeting CCL20-CCR6 signaling not only enhances therapeutic efficacy but also remodels the TME by reducing macrophage infiltration and altering immune cell composition.

### Triple combination therapy induces robust cytotoxicity in glioblastoma organoids

To further evaluate treatment efficacy in a more physiologically relevant setting, we generated GBM organoids (GBOs) from orthotopically implanted CT-2A and GL261 tumors in C57BL/6 mice (**Fig 4A**). These tumors exhibited elevated CCL20/CCR6 expression compared to adjacent tissue and robust macrophage infiltration, as evidenced by IBA1 staining, recapitulating key features observed in human samples (**Fig S5**). Tumors were dissected and cultured under low attachment conditions, as described^45^. CT-2A- and GL261-derived organoids were collected at one- and two-week time points and subjected to transcriptomic analysis by RNA sequencing (RNA-seq). The proportions of mapped reads and overall transcript abundance distributions were consistent across samples. (**Fig S6 A-B**). Organoid transcriptomes were then compared with matched snap-frozen tumor tissue collected at the time of dissection. Principle component analysis (PCA) of normalized gene expression values revealed distinct clustering by time point, indicating temporal changes in gene expression during organoid culture (**Fig 4B**). Despite these changes, Pearson correlation analysis demonstrated strong concordance between organoids and their corresponding primary tumors (r = 0.75-0.9), underscoring the preservation of global transcriptomic features (**Fig 4C, S6C**). Notably, CCL20 expression remained stable in both CT-2A and GL261 organoids at one and two weeks, indicating maintenance of *in vivo* expression levels.

**Figure 4.**
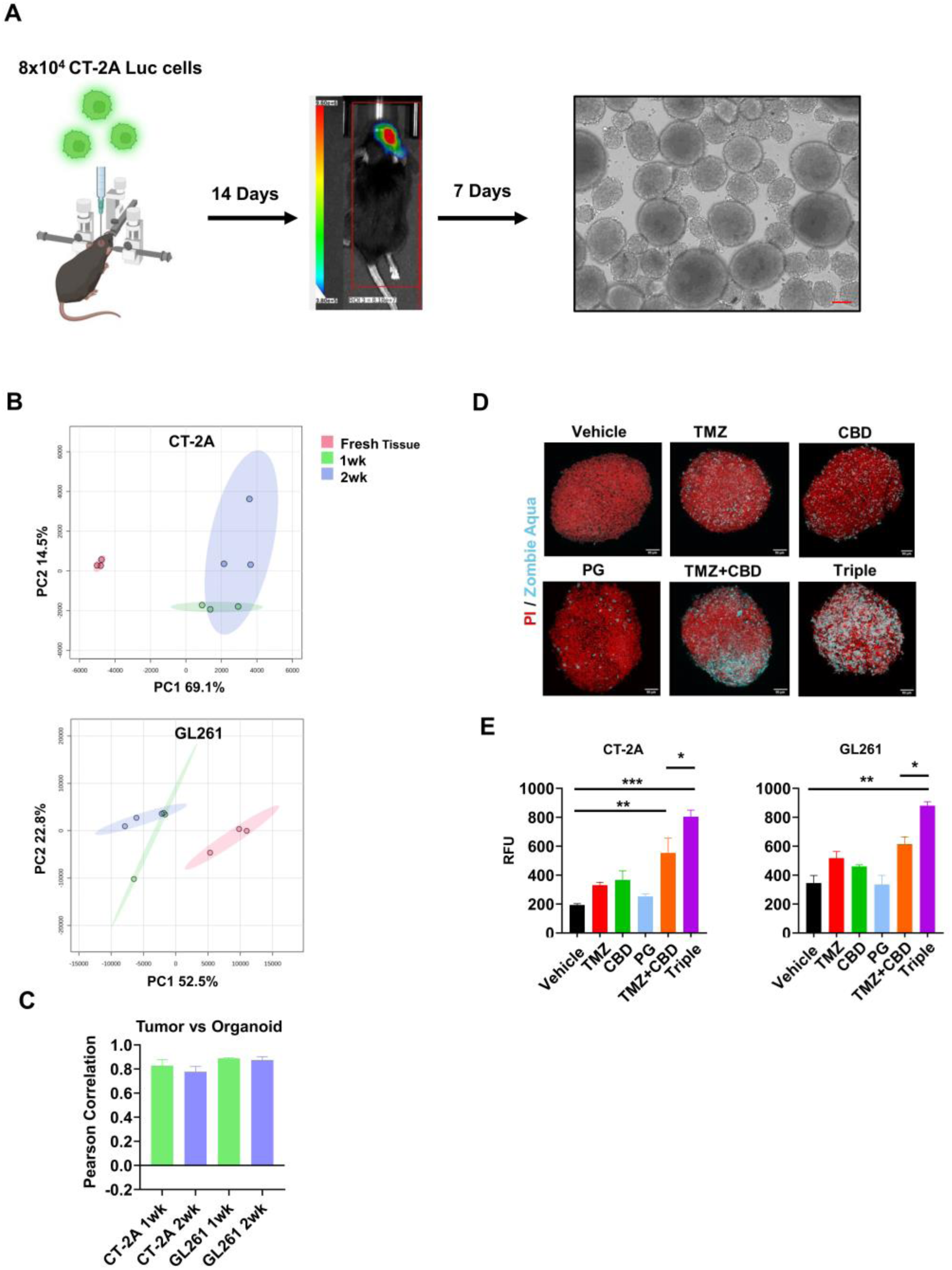
Inhibition of CCL20/CCR6 enhances TMZ + CBD combination therapy in GBM organoids. The effectiveness of single-agent and combination treatments were evaluated ex vivo using an orthotopic GBM organoid model. **(A)** Schematic depicting orthotopic tumor growth and subsequent organoid generation. **(B)** GBM organoid cultures were established from brain tumors from mice implanted with CT-2A or GL261 cells. Poly(A)+ RNA sequencing was performed on both fresh tumor tissue and organoid samples collected after 1 or 2 weeks in culture. PCA of normalized aligned counts (FPKM) was used to assess similarity between samples and groups. **(C)** Pearson correlation was performed to quantify gene expression similarity across tumor and organoid samples. **(D)** Organoids were treated with dug combinations including TMZ, CBD, and PG, and stained with the cell death indicator Zombie Aqua (cyan) and PI (red). Organoid images were captured as z-stacks using laser scanning confocal microscopy (Olympus FV1200). **(E)** Relative zombie aqua signal intensity was quantified from z-projection images of CT-2A and GL261 organoids using ImageJ. Statistical analysis was performed using one-way ANOVA with Holm Sidack correction (*=p≤0.05, **=p≤0.01, ***=p≤0.001).

Drug screening in GBOs presents inherent technical challenges, including variability in organoid size, heterogeneity in organoid number per well, and the suspension-based culture format. To address these limitations, we employed a fixable fluorescent viability dye (Zombie Aqua) in combination with confocal microscopy to quantify treatment-induced cytotoxicity. On day 15 of culture, GBOs were transferred to low-attachment 96-well plates (5-6 organoids per well) and treated with CBD (10 µM), TMZ (500 µM), or the CCL20-modulating agent pioglitazone (PG) (10 µM), either as single agents or in combination, alongside vehicle controls. PG was selected based on prior reports demonstrating its ability to downregulate CCL20 expression^46^. CT2A-derived organoids showed limited sensitivity to single-agent treatments at the tested doses. Modest increases in Zombie Aqua staining were observed in the CBD-, TMZ-, and PG-treated groups relative to vehicle controls, indicating slight increases in membrane permeability and cell death; however, these effects were not statistically significant. In contrast, combination treatments exhibited significantly greater cytotoxic effects. Both the TMZ+CBD and triple combination (TMZ+CBD+PG) groups exhibited significantly increased Zombie staining, with the triple combination eliciting the strongest signal, consistent with enhanced cell death (**Fig 4D**). A similar trend was observed in GL261-derived organoids, where the triple combination treatment resulted in significantly higher cytotoxicity compared to both vehicle and TMZ+CBD treatments (**Fig 4E**). Collectively, these findings demonstrate that modulation of CCL20 signaling enhances the cytotoxic effects of TMZ and CBD in 3D GBM models, supporting its role in therapeutic resistance and highlighting its potential as a combinatorial therapeutic target.

### Integrated proteomic analysis uncovers immune, metabolic, and mitochondrial pathways mediating triple-combination efficacy

We performed mass spectrometry-based proteomic profiling of treated tumors (as described Fig. 3A) to define the mechanisms underlying the synergy between TMZ, CBD, and CCL20 inhibition. Differential expression analysis revealed substantial, treatment-dependent remodeling of the tumor proteome with 2,195 differentially expressed proteins (DEPs) in CBD, 588 in TMZ, 44 in shDPX, 727 in TMZ+CBD, and 1,140 in the triple combination versus vehicle, along with 135 DEPs in triple combination versus TMZ+CBD. The uniqueness and overlap of DEPs across treatment groups are illustrated by Circos plot (**Fig 5A**), while global patterns of protein regulation are visualized using volcano plots (**Fig 5B**). Although no DEPs were shared across all treatment groups, partial convergences between shDPX and CBD (25 shared DEPs) and substantial overlap between the triple combination and TMZ (312 DEPs) or CBD (403 DEPs), were observed indicating that the triple regimen integrates and extends single-agent effects.

**Figure 5.**
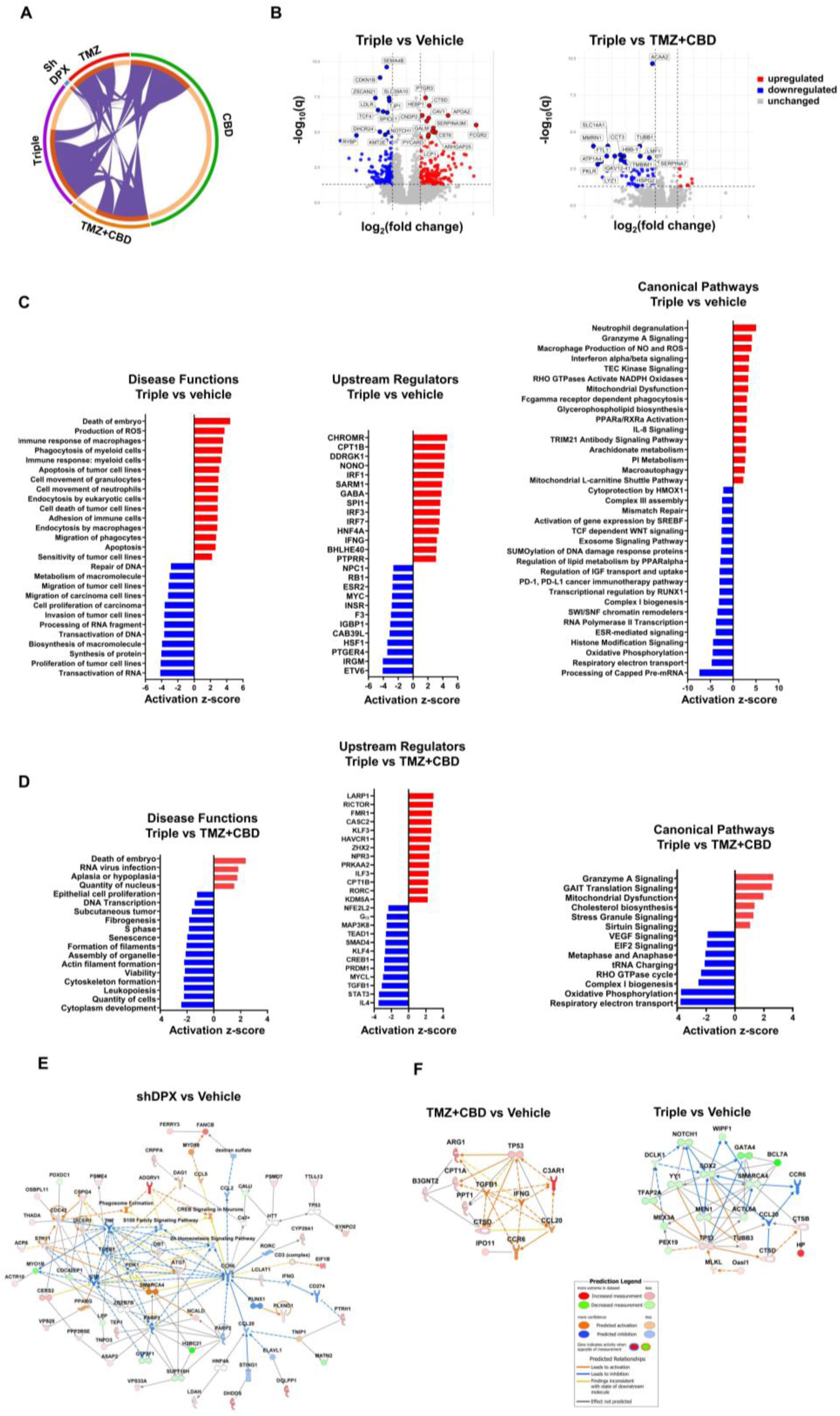
Proteomic analysis reveals functional consequences of CCL20 inhibition. CT-2A tumors were collected and subjected to mass spectrometry-based proteomic analysis. **(A)** A circos plot illustrating the overlap of DEPs across treatment groups. Outer arcs represent individual DEP datasets, color-coded according to the treatment groups in figure 3A. Inner arcs depict gene members within each dataset. Dark orange denotes proteins shared across multiple datasets, whereas light orange color indicates proteins unique to a given list. Purple connecting lines link shared proteins across datasets. **(B)** Volcano plots showing DEPs for comparisons of TMZ+CBD vs Vehicle, Triple combination vs Vehicle, and Triple combination vs TMZ+CBD. **(C-D)** Functional pathway analyses of DEPs were performed using IPA for (C) Triple combination vs Vehicle and (D) Triple combination vs TMZ+CBD comparisons. Functional enrichment was calculated using the right-tailed Fisher’s exact test. The top activated and inhibited disease functions, upstream regulators, and canonical pathways are shown, ranked by their activation z-score. **(E)** Protein interaction network generated from DEPs identified in the shDPX vs vehicle comparison, highlighting connections to CCR6 signaling. **(F)** Protein interaction networks were centered on CCL20 were constructed using DEPs (p≤0.05) from either the TMZ+CBD vs Vehicle or the Triple vs Vehicle comparisons. Activation or inhibition of CCL20 signaling was predicted using IPA.

Ingenuity Pathway Analysis (IPA **Fig. 5C**) revealed that the triple combination activates phagocytosis, ROS production, myeloid immune responses, and apoptosis, while suppressing RNA transcription, protein and macromolecule synthesis, and tumor proliferation. Key upstream regulators included the cholesterol-induced regulator of metabolism RNA (CHROMR), carnitine palmitoyl-transferase 1B (CPT 1B), ER-stress induced DDRGK1, and the prostaglandin E receptor 4. These changes were accompanied by activation of immune pathways (neutrophil degranulation, granzyme A and interferon signaling) and metabolic disruptions (mitochondrial dysfunction, PPARα activation, and phospholipid metabolism), alongside suppression of oxidative phosphorylation and electron transport chain activity.

Comparison with TMZ+CBD (**Fig 5D**) highlighted the specific impact of CCL20 inhibition, revealing reduced tumor growth, viability, and cell number, along with prominent cell cycle disruption and suppressed S-phase progression (**Fig S7A**). Predicted regulators included LARP1, RICTOR, TGFB1, STAT3, and IL4. This condition was marked by enhanced mitochondrial stress, reduced complex I biogenesis, and further suppression of oxidative phosphorylation, as well as activation of Granzyme A signaling and cholesterol biosynthesis.

shDPX monotherapy induced limited proteomic changes but activated lipid synthesis, autophagy, and anti-tumor immune responses while suppressing metastasis and neoplasia **(Fig S7B)**. In contrast, TMZ+CBD drove broad immune activation and metabolic remodeling, including increased lipid catabolism, but also engaged compensatory survival pathways that may limit therapeutic efficacy.

Finally, network analysis confirmed effective suppression of CCL20 signaling (**Fig 5E-F**). While CCL20 itself was not differentially expressed, shDPX reduced CCR6 activity (**Fig 5E**), and TMZ+CBD activated CCL20 signaling (via TGFβ and IFNγ) (**Fig 5F).** Notably, the triple combination suppressed CCL20/CCR6 axis, with key contributions from CTSD and SOX2, supporting effective suppression of the pathway upon shDPX treatment.

### CCL20 inhibition reprograms tumor metabolism and activates 17-HDHA–associated cytotoxic pathways

Given the prominent alterations in lipid metabolic pathways identified in the proteomic analysis, we next performed targeted lipidomic profiling to assess treatment-dependent changes in bioactive lipids. A panel of 55 structurally and functionally diverse lipids was quantified from CT-2A flank tumor extracts using a triple quadrupole mass spectrometer. Partial Least Squares Discriminant Analysis (PLS-DA) revealed separation of treatment groups along the first two principal components, with distinct clustering patterns (**Fig 6A**). Hierarchical clustering (**Fig 6B**) with heatmap visualization further demonstrated treatment-specific lipidomic signatures, indicating broad remodeling of lipid metabolism across experimental conditions (**Fig 6B**). Although the relative abundance of proinflammatory mediators, such as Hydroxyeicosatetraenoic acid (HETEs), and pro-resolving mediators, including Hydroxyeicosapentaenoic acids (HEPEs), and Epoxyeicosatrienoic acids (EETs), remained largely unchanged (**Fig S8**), the triple-combination therapy significantly increased levels of 14- and 17-Hydroxydocosahexaenoic Acid (17-HDHA) compared to TMZ alone (**Fig 6C**).

**Figure 6.**
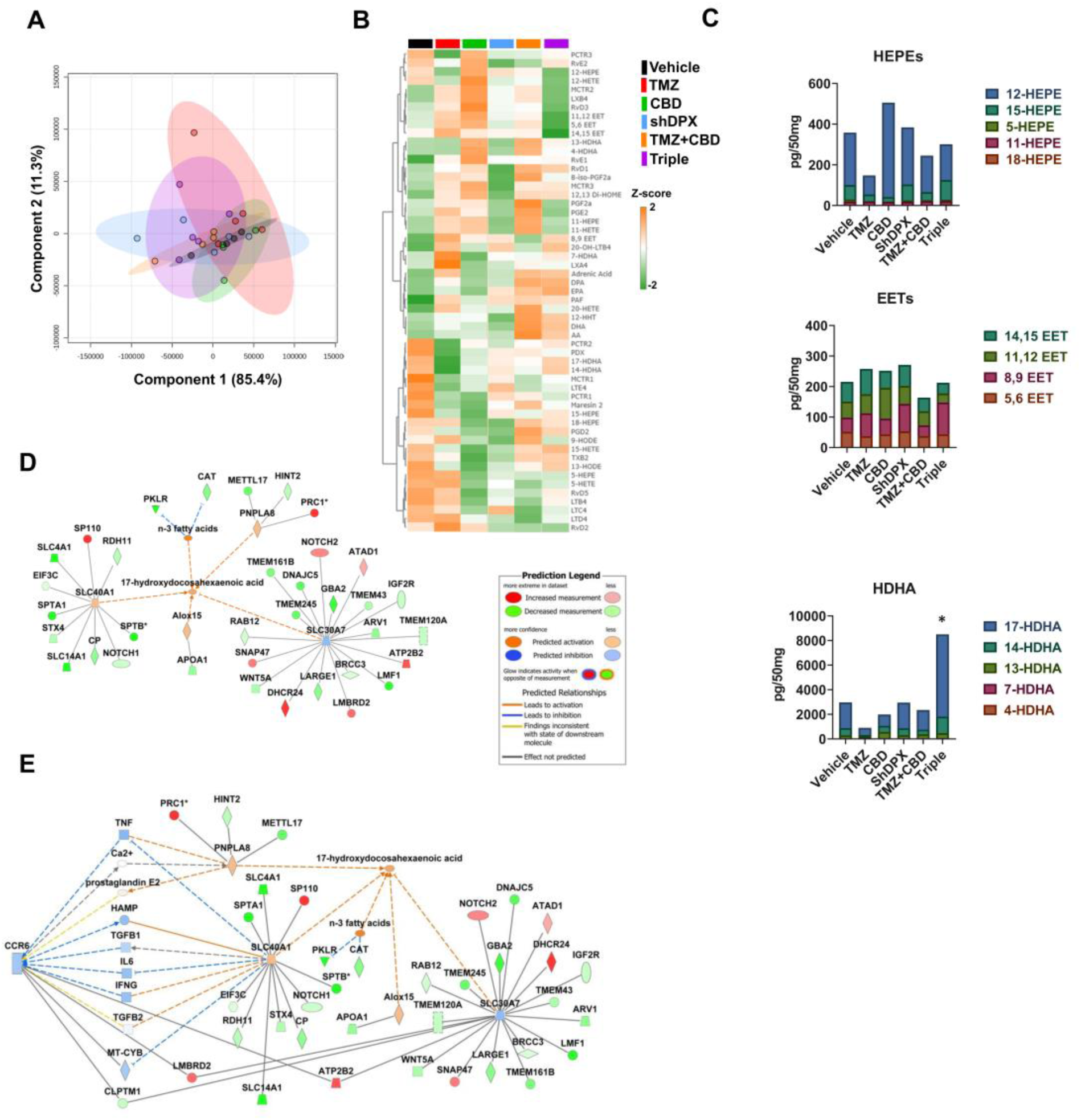
CCL20/CCR6 inhibition reprograms lipid metabolism in GBM tumors. Treated subcutaneous CT-2A tumors were analyzed for the abundance of a panel of bioactive lipids by LC-MS. **(A)** PLS-DA showing supervised clustering of tumor samples based on lipid abundance across treatment groups, visualized using the first two PLS components (X and Y axes). **(B)** Hierarchical clustering of z-score-normalized lipid abundance across treatment groups. **(C)** Intratumoral levels of pro-resolving lipid mediators stratified by treatment group. (n=5, one-way ANOVA with Holm Sidak correction *=p≤0.05 vs vehicle). **(D)** Protein-lipid interaction network was generated using IPA to investigate the interactions between DEPs (p≤0.05) from the Triple combination vs TMZ+CBD dataset and 17-HDHA. **(E)** Network analysis illustrating that inhibition of CCR6 signaling is associated with increased 17-HDHA levels. Upregulated (red) and downregulated (green) intermediary proteins identified in the Triple combination vs TMZ+CBD DEP dataset are shown.

To further investigate this finding, we examined interactions between 17-HDHA and the DEPs identified in the proteomic dataset. IPA-based protein-lipid interaction network modelling revealed strong concordance between the lipidomic and proteomic data, with 17-HDHA predicted to be activated based on DEP fold changes (**Fig 6D**). This predicted activation was primarily driven by SLC40A1, SLC30A7, which function as iron and zinc transporters, respectively, as well as PNPLA8, a calcium-independent phospholipase linked to mitochondrial function. Notably, DHA and specifically its conversion to 17-HDHA via the intermediate 170HpDHA has been reported to induce tumor cell toxicity in part through increased ROS production^47^. To further define the role of CCL20 in regulating lipid metabolism in this context, we mapped interactions between CCR6 and the 17-HDHA regulatory network. This analysis identified several key signaling mediators, including TNF, IL6, TGFβ, IFN-γ, the mitochondrial protein cytochrome B, and the lipid scramblase CLPTM1 (**Fig6 E**). Collectively, these findings demonstrate that inhibition of CCL20 signaling promotes metabolic reprogramming characterized by selective enrichment of 17-HDHA and engagement of lipid-associated cytotoxic pathways, providing mechanistic insight into the enhanced efficacy of the triple-combination therapy.

### Triple combination selectively amplifies mitochondrial superoxide generation

With both proteomic and lipidomic datasets converging on mitochondrial metabolism and ROS production as central features, we next sought to characterize the impact of CCL20/CCR6 inhibition on mitochondrial dysfunction. To this end, we examined protein expression changes uniquely associated with the triple combination. Given that CCL20 signaling was activated in the TMZ+CBD condition but suppressed in the triple combination, we compared differential expression networks to identify proteins exhibiting treatment-specific patterns. Specifically, we focused on proteins uniquely regulated in the triple combination (i.e., upregulated in triple combination vs. TMZ+CBD while downregulated in TMZ+CBD vs. vehicle, or vice versa). Interrogation of these proteins using IPA revealed key intermediates linking CCR6 signaling to mitochondrial dysfunction, including SLC4A2, TTYH3, ATP2B2, NDUFA1, and CYB5R3 (**Fig. 7A**). These findings suggest that CCL20 inhibition exacerbates mitochondrial dysfunction beyond that induced by TMZ+CBD alone.

**Figure 7.**
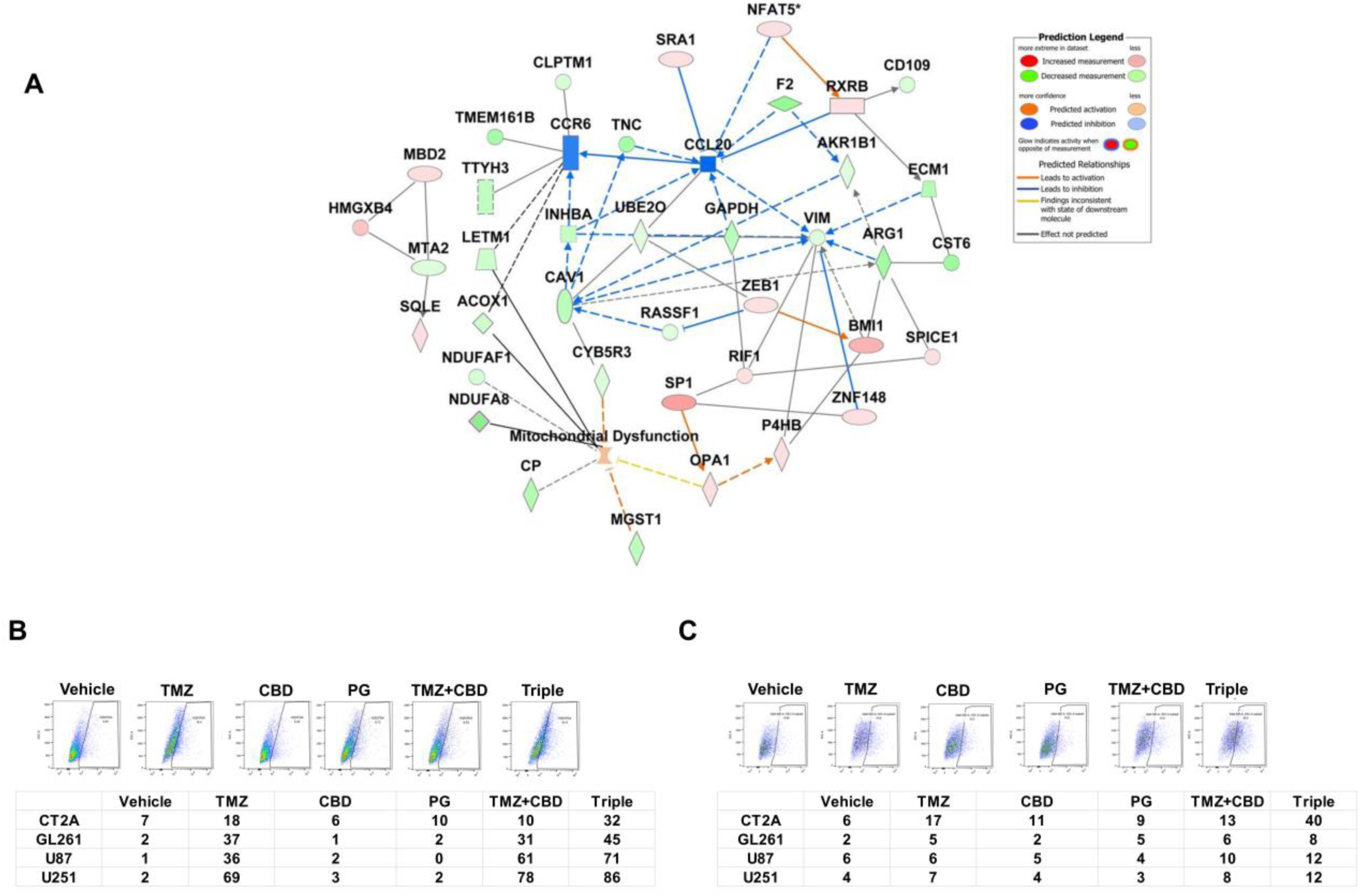
TMZ, CBD and CCL20 inhibition converge on mitochondrial stress and ROS generation. Effects specific to CCL20 inhibition were investigated using the differential expression dataset comparing triple combination vs TMZ+CBD treatment. **(A)** Network analysis of DEPs uniquely associated with the triple combination reveals key intermediaries linking CCR6 signaling to mitochondrial dysfunction. **(B-C)** GBM cell lines were treated with PG (10µM), CBD (10µM), TMZ (500µM), or the indicated combinations. (B) Total cellular ROS levels were measured by CM-H2DCFDA staining and quantified by flow cytometry. and (C) Mitochondrial superoxide levels were evaluated using MitoSox red staining followed by flow cytometry. Representative dot plots are shown along with percentage of positive cells for each cell line and treatment condition.

To functionally validate these observations, we quantified total cellular ROS and mitochondrial-specific ROS following treatment with individual agents or their combinations. CT-2A, GL261, U87 and U251 cells were treated for 48 hours with TMZ, CBD, PG, double combinations, or the triple combination. Cells were then stained with the pan-ROS indicator CM-H2DCFDA or the mitochondrial superoxide-specific probe MitoSox Red, and fluorescence intensity was quantified using flow cytometry (**Fig 7B-C**). TMZ monotherapy markedly increased total ROS levels across all cell lines. However, in most cases, the CM-H2DCFDA signal intensity in the TMZ+CBD group was comparable to or lower than TMZ alone. In contrast, the triple combination consistently induced greater increase in total ROS levels than TMZ

## Discussion

The present study identifies the CCL20-CCR6 axis as a critical mediator of therapeutic resistance in GBM and demonstrates that its inhibition enhances the efficacy of TMZ, alone or in combination with CBD. CCL20 is elevated in GBM, correlates with poor patient survival, and is further induced by TMZ±CBD, suggesting a stress-responsive, pro-survival role. Collectively, these findings position CCL20 as both a potential predictive biomarker of recurrence and treatment response, and a promising therapeutic target for improving outcomes in TMZ-treated GBM.

Consistent with prior reports, we confirm the prognostic relevance of CCL20 in glioma and GBM patients using multiple publicly available datasets and extend this observation by showing that its association with survival is specific to TMZ-treated patients, but not those receiving bevacizumab. This treatment-specific effect highlights its potential clinical utility.

Another key finding of this study is the robust synergy between TMZ and CBD across murine and human GBM models, including those with diverse oncogenic backgrounds. However, both treatments induce CCL20, indicating activation of adaptive resistance. Pharmacologic or genetic disruption of CCL20/CCR6 significantly enhances TMZ+CBD efficacy, both *in vivo* and in organoid models, establishing CCL20 as a key compensatory mechanism limiting response.

Mechanistically, multi-omics analyses reveal that combined TMZ, CBD, and CCL20 inhibition converge on mitochondrial dysfunction, metabolic reprogramming, and oxidative stress (**Fig 8**). Mitochondrial function is critical for GBM growth, as oxidative respiration supports pyrimidine synthesis^48^. Accordingly, disrupting mitochondrial respiration may impair DNA repair by limiting nucleotide availability. GBM cells can partially compensate via horizontal mitochondrial transfer from astrocytes, microglia, and neurons^49,50^, highlighting both the importance of mitochondrial integrity and a potential therapeutic vulnerability. TMZ induces DNA damage-associated ROS ^51,52^ and mitochondrial dysfunction^53^. Our data extends this model by implicating CCL20 as a regulatory component of the stress response. Specifically, CCL20 appears to act as a buffering node that limits amplification of TMZ-induced oxidative damage, although further studies are needed to fully define this mechanism. Inhibition of CCL20 removes this constraint, promoting a feed-forward cycle of DNA damage and ROS accumulation that enhances cytotoxicity.

**Figure 8.**
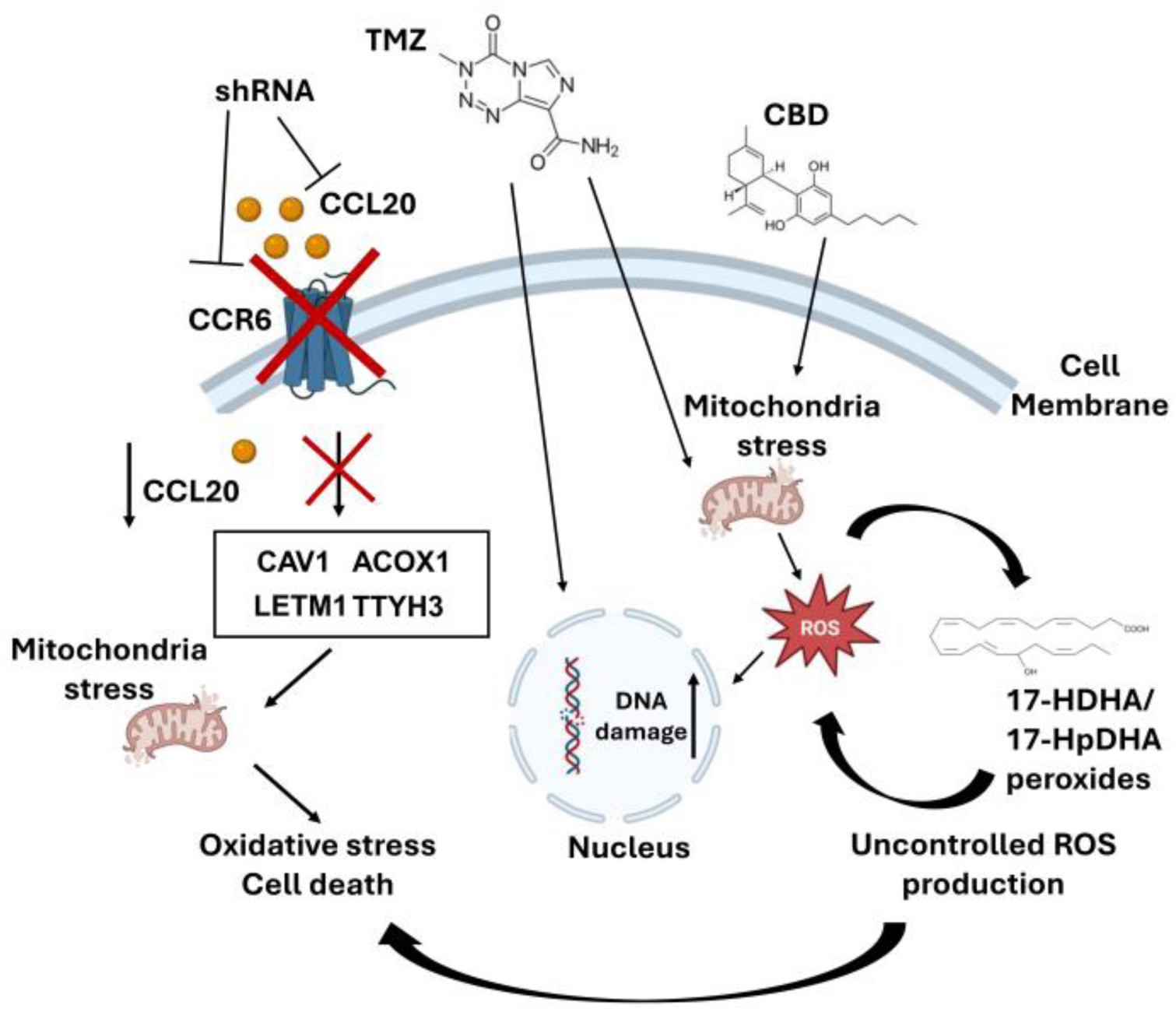
Summary of the role of CCL20/CCR6 signaling in TMZ+CBD treatment response. Graphical summary illustrating of convergent effects of TMZ, CBD, and CCL20 inhibition driving mitochondrial dysfunction, oxidative stress, and tumor cell death.

Proteomic analysis identified key downstream mediators of CCR6 signaling that may link this pathway to mitochondrial homeostasis, including proteins involved in mitochondrial lipid metabolism such as caveolin-1, Tweety homolog 3, and acyl-CoA oxidase 1^54–56^. Consistent with this model, the triple combination uniquely amplifies mitochondrial oxidative stress. While TMZ alone increased total ROS, its effect on mitochondrial superoxide was modest. In contrast, addition of CBD and CCL20 inhibition markedly increased mitochondrial ROS, particularly in immunocompetent CT-2A cells. Together, these findings highlight mitochondrial ROS as a key functional output of the triple therapy and suggest that mitochondrial vulnerability underlies the observed therapeutic synergy.

Beyond tumor cell-intrinsic effects, our proteomic analysis indicates that the triple combination also remodels the immune microenvironment, enhancing phagocyte activity, macrophage-associated ROS production, and granzyme-A signaling. CBD likely contributes substantially to these effects, as it has been shown to modulate the glioma TME through regulation of IL-8, indoleamine 2,3-dioxygenase pathways, and to promote T-cell proliferation either alone or in combination with other immunomodulatory agents^57,58^. Consistent with these findings, immunophenotyping of splenocytes from tumor-bearing mice revealed a broad increase in systemic CD8^+^ T cells and reduced M-MDSCs, consistent with reduced immunosuppression. While CBD has been reported to suppress MDSC proliferation in colorectal adenomas^59^, CCL20 is well known to recruit MDSCs to brain tumors^60^, suggesting that combined targeting of these pathways may synergistically remodel the immunosuppressive TME.

Our results also implicate CCL20/CCR6 signaling in metabolic adaptation. While CCR6 signaling is linked to PI3K-Akt-mTOR and STAT3 pathways^61,62^, TMZ+CBD treatment activated lipid catabolism pathways that were suppressed by CCL20 inhibition. Additionally, convergence of CBD-induced calcium dysregulation with CCR6-associated signaling may further amplify mitochondrial stress and ROS production^63^.

Lipidomic analyses further reinforce the role of mitochondrial oxidative stress and reveal an additional layer of metabolic remodeling associated with treatment response. Notably, the triple combination selectively increased levels of 17-HDHA, a bioactive derivative of DHA^64,65^. Integration of lipidomic and proteomic datasets demonstrated concordant activation of 17-HDHA-associated networks, driven in part by proteins involved in metal ion transport (SLC40A1, SLC30A7) and mitochondrial lipid metabolism (PNPLA8). These findings suggest that altered lipid handling and redox-active metabolite production contribute to treatment efficacy. The biological role of 17-HDHA in this context is likely multifaceted. While often described as a pro-resolving mediator with protective effects in non-malignant tissues, its biosynthetic intermediate, 17-hydroperoxy-DHA (17-HpDHA), has been shown to exhibit cytotoxic activity through lipid peroxidation and ROS generation^47^. The lack of downstream resolvin production suggests accumulation of cytotoxic lipid intermediates that may further drive tumor cell death. Given that lipid peroxidation products can both result from and propagate ROS, this may represent an additional feed-forward mechanism contributing to tumor cell death.

These findings further elucidate the contribution of CBD to the efficacy of the triple combination therapy in GBM. CBD is known to induce oxidative stress in cancer cells, it can also exert antioxidant effects in non-malignant contexts, suggesting a degree of tumor selectivity that may enhance the therapeutic window of combination treatments^66,67^. Moreover, CBD is known to modulate lipid metabolism and shift polyunsaturated fatty acid balance toward n-3 species, potentially facilitating increased production of DHA-derived metabolites such as 17-HDHA^68^. Collectively, these observations indicate that CBD likely plays a dual role in mediating the therapeutic effects of the combination regimen.

Collectively, these findings support a model in which TMZ induces CCL20-dependent survival signaling, CBD amplifies metabolic and redox stress, and CCL20 inhibition of removes a key resistance node, resulting in sustained mitochondrial dysfunction, lipid peroxidation, and ROS-driven cytotoxicity.

These findings have important translational implications. Targeting mitochondrial metabolism and redox homeostasis represents an emerging strategy in GBM, and our data suggest that combining TMZ with CBD and CCL20/CCR6 inhibitors may provide synergistic benefit. Notably, in our model these three regimens were administered separately, however, the identification of safe and effective small molecule inhibitors of CCL20-CCR6 axis may enable the development of an oral combination therapy for TMZ-resistant GBM. In addition to therapeutic applications, components of this pathway, including CCL20 expression and lipid mediators such as 17-HDHA, may serve as biomarkers for patient stratification and response monitoring.

Future studies should explore the potential involvement of PPARγ signaling, given that CBD, PG, and 17-HDHA are all known activators of this pathway^64,69,70^, and further define the immunological and metabolic consequences of CCL20 inhibition. Elucidating these interactions may reveal additional therapeutic strategies to overcome TMZ resistance in GBM.

## Methods

### Bioinformatics

Aligned RNA-seq data were obtained from the NIH Genomic Data Commons (GDC; dbGaP access). Cohorts included untreated primary or recurrent glioma solid tissue samples. Alignment was performed using the GDC standard pipeline. Differential gene expression analysis was conducted in R (DESeq2), with significance defined as adjusted *p* ≤ 0.05 (Benjamini–Hochberg) and log2 fold change calculated. Survival data were generated using the GDC cohort builder with specified treatment stratifications (temozolomide ± bevacizumab). PCA plots of FPKM matrices were generated using MetaboAnalyst. RNA-seq and statistical analyses were performed by LC Sciences.

### Cell Culture

Mouse (CT2A, GL261, SMA-560) and human (U87, U251) glioma cell lines were maintained in DMEM supplemented with 10% FBS and 1% antibiotics at 37°C in 5% CO₂.

### Organoid Culture

Orthotopic tumors were established in C57BL/6 mice by intracranial injection of 8×10 CT2A-luc or GL261-luc cells. Tumor growth was monitored by IVIS imaging. At 2 weeks, tumors were harvested, sectioned (2–5 mm), and cultured in low-attachment plates with organoid media on an orbital shaker (120 rpm, 37°C, 5% CO₂). At day 15, organoids were treated with CBD (10 µM), TMZ (500 µM), pioglitazone (10 µM), combinations, or vehicle. After 48 h, samples were stained (Zombie Aqua, PI), fixed, and imaged by confocal microscopy.

### CRISPR/Cas9 Editing

CT2A cells were transfected with three all-in-one CRISPR plasmids targeting CCL20 (Genecopoia) using Lipofectamine 3000. mCherry-positive cells were sorted and clonally expanded. Gene editing was confirmed by Sanger sequencing.

### Immunostaining and Imaging

Tissue sections (30 µm) and microarrays underwent antigen retrieval, permeabilization, and blocking before overnight primary antibody incubation. Fluorescent or biotinylated secondary antibodies were applied, followed by imaging (Olympus microscopy). For immunoperoxidase staining, sections were treated with hydrogen peroxide, incubated with ABC reagent, and developed with DAB. Images were acquired under identical settings and quantified using ImageJ.

### qRT-PCR

RNA was isolated (Qiagen RNeasy), and cDNA was synthesized (Thermo Maxima kit). Quantitative PCR was performed using SYBR Green on a Bio-Rad CFX384 system. Data were analyzed using CFX Maestro with significance set at *p* ≤ 0.05.

### Cell Viability and Drug Synergy

Cells were plated (5,000 cells/well) and treated with serial dilutions of CBD, TMZ, pioglitazone, or combinations. After 48 h, viability was measured using CellTiter-Glo. Dose–response curves were generated (GraphPad Prism), and drug synergy was assessed using CompuSyn (Chou– Talalay method).

### Dendriplex (shDPX) Preparation

Generation 4 PAMAM dendrimers were sonicated and extruded to ∼5 nm (PDI ∼0.15), then complexed with shRNAs targeting CCL20 and CCR6 to form dendriplex nanoparticles (shDPX), as previously described^44^.

### In Vivo Tumor Studies

C57BL/6 mice were subcutaneously injected with 1×10 CT2A-luc cells. Once tumors reached ∼6 mm, mice received vehicle, CBD (25 mg/kg, oral), TMZ (20 mg/kg, IP), or combination therapy every other day. shDPX or control dendriplex was administered IP twice weekly. Tumors were monitored by caliper, and mice were euthanized on day 30.

### Proteomics

Tumor tissues (n=4/group) were processed using the iST kit and analyzed by diaPASEF (timsTOF Pro). Data were processed with DIA-NN and MS-DAP; differential protein abundance was determined using MSqRob (*p* ≤ 0.01). Pathway analysis was performed using Ingenuity Pathway Analysis, and overlaps visualized via circos plot.

### Lipidomics

Lipids were extracted from tumor tissues and analyzed by targeted LC-MS/MS (Sciex QTRAP 6500+). Data were processed with Analyst and Sciex OS software. Statistical analysis and visualization (heatmaps, PLS-DA) were performed using MetaboAnalyst.

### Flow Cytometry

For ROS analysis, treated cells were stained with CM-H2DCFDA (total ROS) or MitoSOX Red (mitochondrial ROS), with DAPI for viability, and analyzed on a BD LSRII. For immunophenotyping, splenocytes were isolated, RBC-lysed, stained with T cell or myeloid panels, and analyzed by flow cytometry. Data were processed using FlowJo.

monotherapy (**Fig 7B**). Importantly, while TMZ treatment resulted in only modest increases in mitochondrial superoxide, the triple combination elicited a substantially stronger induction in mitochondrial ROS. This effect was particularly pronounced in CT-2A cells (**Fig 7C**), indicating that CCL20 inhibition selectively amplifies mitochondrial superoxide generation. Collectively, these results support a model in which targeting CCL20/CCR6 signaling enhances mitochondrial dysfunction and ROS-mediated cytotoxicity, providing a mechanistic basis for the superior efficacy of the triple-combination therapy.

## Acknowledgement

We would like to acknowledge the USF COM Fred Wright Jr Flow Cytometry Core, the Lisa Muma Weitz Laboratory for Advanced Microscopy & Cell Imaging, the USF Advanced Research Core for Mass Spectrometry (ARC-MS), and the USF Heart Institute Core Facilities for their contributions to this work.

## Author Contributions

Subhra Mohapatra, Shyam Mohapatra, and Ryan Green designed experiments. Ryan Green, Karthick Mayilsamy, Erin Anglin, Kristina Tosi, Sashank Bikkasani, and Parthvi Bharatkumar Patel performed experiments. Eleni Markoutsa prepared dendriplexes, Tiara Wolf, Jennifer Guergues, and Stanley M Stevens performed proteomic analysis. Ganesh Haladae performed lipidomic analysis. Ryan Green drafted the manuscript. Subhra Mohapatra, Shyam Mohapatra, and Ryan Green edited the final draft.

## Conflict of Interest Statement

The authors have no conflicts of interest to declare.

## Funding Statement

This work was supported by a Veterans Affairs Merit Review grant BX005757 awarded to SM, and by Research Career Scientist Awards, IK6BX004212 to SM and IK6BX006032 to SSM. Although this report is based in part on work supported by the Department of Veterans Affairs, Veterans Health Administration, Office of Research and Development, the contents do not represent the views of the Department of Veterans Affairs or the United States Government.

## Data Availability Statement

All data supporting the findings of this study are available within the article including figures and supplementary information. Sequencing data from patient GBM samples can be accessed within the NIH Genomic Data Commons. Sequencing data from organoid samples can be accessed within the NCBI Gene Expression Omnibus database (Accession# GSE-TBD)

## Animal Ethics Statement

All protocols in this study were approved by the University of South Florida Institutional Animal Care and Use Committee, protocol number IS00011567, PHS Assurance number: D-16-00589 (A4100-01), AAALACi Accreditation number: 434, in accordance with the Guide for the Care and Use of Laboratory Animals (Guide), the Animal Welfare Act, the Animal Welfare Regulations (Title 9 Code of Federal Regulations Subchapter A, “Animal Welfare”, Parts 1-3 [AWA]), the Public Health Service Policy on Humane Care and Use of Laboratory Animals (PHS Policy), and University Policy #0-308.

## Notes

### Competing Interest Statement

The authors have declared no competing interest.

