## Supplemental Information for "CCL20–CCR6 Signaling as a Prognostic Biomarker and Therapeutic Target in Temozolomide-Resistant Glioblastoma"

#### Supplemental Information

**Supplementary Table 1. Comparison of Oncogenic Driver Mutations in Mouse and Human GBM Models**

| | MGMT | IDH | P53 | Ras | PTEN | TERT | NF1 |
|---|---|---|---|---|---|---|---|
| CT-2A | - | WT | WT | Nras Q61>L | deficiency | - | - |
| GL261 | Unmethylated | WT | R>P codon 153 of exon 5 | Kras G12>C | deficiency | - | - |
| U87 | Methylated | WT | WT | - | 209G>T | 228C>T | deletion (c3739-3742) |
| U251 | Methylated | WT | R273>H | - | G242>V | C228>T | - |

### Supplementary Fig. 1

| Project ID | Sample ID | Tissue Type | Tumor Descriptor | Specimen Type | Preservation Method |
|---|---|---|---|---|---|
| TCGA-GBM | TCGA-12-3648-01A | Tumor | Primary | Solid Tissue | Unknown |
| TCGA-GBM | TCGA-19-1390-01A | Tumor | Primary | Solid Tissue | Unknown |
| TCGA-GBM | TCGA-19-2620-01A | Tumor | Primary | Solid Tissue | Unknown |
| TCGA-GBM | TCGA-12-3650-01A | Tumor | Primary | Solid Tissue | Unknown |
| TCGA-GBM | TCGA-41-2571-01A | Tumor | Primary | Solid Tissue | Unknown |
| TCGA-GBM | TCGA-02-0026-01B | Tumor | Primary | Solid Tissue | Unknown |
| TCGA-GBM | TCGA-19-1787-01B | Tumor | Primary | Solid Tissue | Unknown |
| TCGA-GBM | TCGA-19-1787-01B | Tumor | Primary | Solid Tissue | Unknown |
| TCGA-GBM | TCGA-32-2632-01A | Tumor | Primary | Solid Tissue | Unknown |
| TCGA-GBM | TCGA-02-0070-01A | Tumor | Primary | Solid Tissue | Unknown |
| TCGA-GBM | TCGA-12-1597-01B | Tumor | Primary | Solid Tissue | Unknown |
| TCGA-GBM | TCGA-32-2615-01A | Tumor | Primary | Solid Tissue | Unknown |
| TCGA-GBM | TCGA-12-3653-01A | Tumor | Primary | Solid Tissue | Unknown |
| TCGA-GBM | TCGA-32-2634-01A | Tumor | Primary | Solid Tissue | Unknown |
| TCGA-GBM | TCGA-12-3652-01A | Tumor | Primary | Solid Tissue | Unknown |
| TCGA-GBM | TCGA-32-2638-01A | Tumor | Primary | Solid Tissue | Unknown |
| TCGA-GBM | TCGA-14-2555-01B | Tumor | Primary | Solid Tissue | Unknown |
| TCGA-GBM | TCGA-12-3652-01A | Tumor | Primary | Solid Tissue | Unknown |
| TCGA-GBM | TCGA-12-3653-01A | Tumor | Primary | Solid Tissue | Unknown |
| TCGA-GBM | TCGA-19-2624-01A | Tumor | Primary | Solid Tissue | Unknown |
| TCGA-GBM | TCGA-19-2619-01A | Tumor | Primary | Solid Tissue | Unknown |
| TCGA-GBM | TCGA-32-2616-01A | Tumor | Primary | Solid Tissue | Unknown |
| TCGA-GBM | TCGA-19-2619-01A | Tumor | Primary | Solid Tissue | Unknown |
| TCGA-GBM | TCGA-02-0016-01A | Tumor | Primary | Solid Tissue | Unknown |
| TCGA-GBM | TCGA-12-3650-01A | Tumor | Primary | Solid Tissue | Unknown |
| TCGA-GBM | TCGA-19-2625-01A | Tumor | Primary | Solid Tissue | Unknown |
| TCGA-GBM | TCGA-19-2629-01A | Tumor | Primary | Solid Tissue | Unknown |
| TCGA-GBM | TCGA-12-3646-01A | Tumor | Primary | Solid Tissue | Unknown |
| TCGA-GBM | TCGA-02-0039-01A | Tumor | Primary | Solid Tissue | Unknown |
| TCGA-GBM | TCGA-12-3651-01A | Tumor | Primary | Solid Tissue | Unknown |
| TCGA-GBM | TCGA-19-2629-01A | Tumor | Primary | Solid Tissue | Unknown |
| TCGA-GBM | TCGA-12-3644-01A | Tumor | Primary | Solid Tissue | Unknown |
| HCMI-CMDC | HCM-BROD-0418-C71-01A | Tumor | Primary | Solid Tissue | Frozen |
| HCMI-CMDC | HCM-BROD-0423-C71-01A | Tumor | Primary | Solid Tissue | Frozen |
| HCMI-CMDC | HCM-BROD-0416-C71-01A | Tumor | Primary | Solid Tissue | Frozen |
| HCMI-CMDC | HCM-BROD-0796-C71-01B | Tumor | Primary | Solid Tissue | Frozen |
| HCMI-CMDC | HCM-BROD-0199-C71-01A | Tumor | Primary | Solid Tissue | Frozen |
| HCMI-CMDC | HCM-BROD-0196-C71-01A | Tumor | Primary | Solid Tissue | Frozen |
| HCMI-CMDC | HCM-BROD-0455-C71-01A | Tumor | Primary | Solid Tissue | Frozen |
| HCMI-CMDC | HCM-BROD-0011-C71-01A | Tumor | Primary | Solid Tissue | Frozen |
| HCMI-CMDC | HCM-BROD-0613-C71-01A | Tumor | Primary | Solid Tissue | Frozen |
| HCMI-CMDC | HCM-BROD-0417-C71-01A | Tumor | Primary | Solid Tissue | Frozen |
| HCMI-CMDC | HCM-BROD-0012-C71-01A | Tumor | Primary | Solid Tissue | Frozen |
| HCMI-CMDC | HCM-BROD-0028-C71-01A | Tumor | Primary | Solid Tissue | Frozen |
| HCMI-CMDC | HCM-BROD-0014-C71-01A | Tumor | Primary | Solid Tissue | Frozen |
| HCMI-CMDC | HCM-BROD-0200-C71-01A | Tumor | Primary | Solid Tissue | Frozen |
| HCMI-CMDC | HCM-BROD-0457-C71-01B | Tumor | Primary | Solid Tissue | FFPE |
| HCMI-CMDC | HCM-BROD-0103-C71-01A | Tumor | Primary | Solid Tissue | Frozen |
| HCMI-CMDC | HCM-BROD-0003-C71-01A | Tumor | Primary | Solid Tissue | Frozen |
| HCMI-CMDC | HCM-BROD-0693-C71-01A | Tumor | Primary | Solid Tissue | FFPE |
| HCMI-CMDC | HCM-BROD-0195-C71-01A | Tumor | Primary | Solid Tissue | Frozen |
| HCMI-CMDC | HCM-BROD-0614-C71-01A | Tumor | Primary | Solid Tissue | Frozen |
| HCMI-CMDC | HCM-BROD-1121-C71-01A | Tumor | Primary | Solid Tissue | Frozen |
| HCMI-CMDC | HCM-BROD-0685-C71-01A | Tumor | Primary | Solid Tissue | Frozen |
| TCGA-GBM | TCGA-06-0190-02A | Tumor | Recurrence | Unknown | Unknown |
| TCGA-GBM | TCGA-19-0957-02A | Tumor | Recurrence | Unknown | Unknown |
| TCGA-GBM | TCGA-19-4065-02A | Tumor | Recurrence | Unknown | Unknown |
| TCGA-GBM | TCGA-06-0221-02A | Tumor | Recurrence | Unknown | Unknown |
| TCGA-GBM | TCGA-06-0210-02A | Tumor | Recurrence | Unknown | Unknown |
| TCGA-GBM | TCGA-19-1389-02A | Tumor | Recurrence | Unknown | Unknown |
| TCGA-GBM | TCGA-06-0125-02A | Tumor | Recurrence | Unknown | Unknown |
| TCGA-GBM | TCGA-14-1034-02B | Tumor | Recurrence | Unknown | Unknown |
| TCGA-GBM | TCGA-06-0171-02A | Tumor | Recurrence | Unknown | Unknown |
| TCGA-GBM | TCGA-06-0211-02A | Tumor | Recurrence | Unknown | Unknown |
| TCGA-GBM | TCGA-06-0152-02A | Tumor | Recurrence | Unknown | Unknown |
| TCGA-GBM | TCGA-14-1402-02A | Tumor | Recurrence | Unknown | Unknown |
| TCGA-GBM | TCGA-14-1402-02A | Tumor | Recurrence | Unknown | Unknown |
| TCGA-GBM | TCGA-14-0736-02A | Tumor | Recurrence | Unknown | Unknown |
| HCMI-CMDC | HCM-BROD-0695-C71-02B | Tumor | Recurrence | Solid Tissue | FFPE |
| HCMI-CMDC | HCM-BROD-0106-C71-02A | Tumor | Recurrence | Solid Tissue | Frozen |
| HCMI-CMDC | HCM-BROD-0046-C71-02A | Tumor | Recurrence | Solid Tissue | Frozen |
| HCMI-CMDC | HCM-BROD-0209-C71-02A | Tumor | Recurrence | Solid Tissue | Frozen |
| HCMI-CMDC | HCM-BROD-0199-C71-02A | Tumor | Recurrence | Solid Tissue | Frozen |
| HCMI-CMDC | HCM-BROD-0047-C71-02A | Tumor | Recurrence | Solid Tissue | Frozen |
| HCMI-CMDC | HCM-BROD-0689-C71-02A | Tumor | Recurrence | Solid Tissue | Frozen |
| HCMI-CMDC | HCM-BROD-0415-C71-02B | Tumor | Recurrence | Solid Tissue | Frozen |
| HCMI-CMDC | HCM-BROD-0013-C71-02A | Tumor | Recurrence | Solid Tissue | Frozen |
| HCMI-CMDC | HCM-BROD-0210-C71-02A | Tumor | Recurrence | Solid Tissue | Frozen |
| HCMI-CMDC | HCM-BROD-0213-C71-02A | Tumor | Recurrence | Solid Tissue | Frozen |
| HCMI-CMDC | HCM-BROD-1123-C71-02A | Tumor | Recurrence | Solid Tissue | Frozen |
| HCMI-CMDC | HCM-BROD-0681-C71-02A | Tumor | Recurrence | Solid Tissue | Frozen |
| HCMI-CMDC | HCM-BROD-0830-C71-02A | Tumor | Recurrence | Solid Tissue | Frozen |

Supplementary Figure 1. List of samples used for differential gene expression analysis between primary and recurrent GBM patients in figure 1D.

#### Supplementary Fig. 2

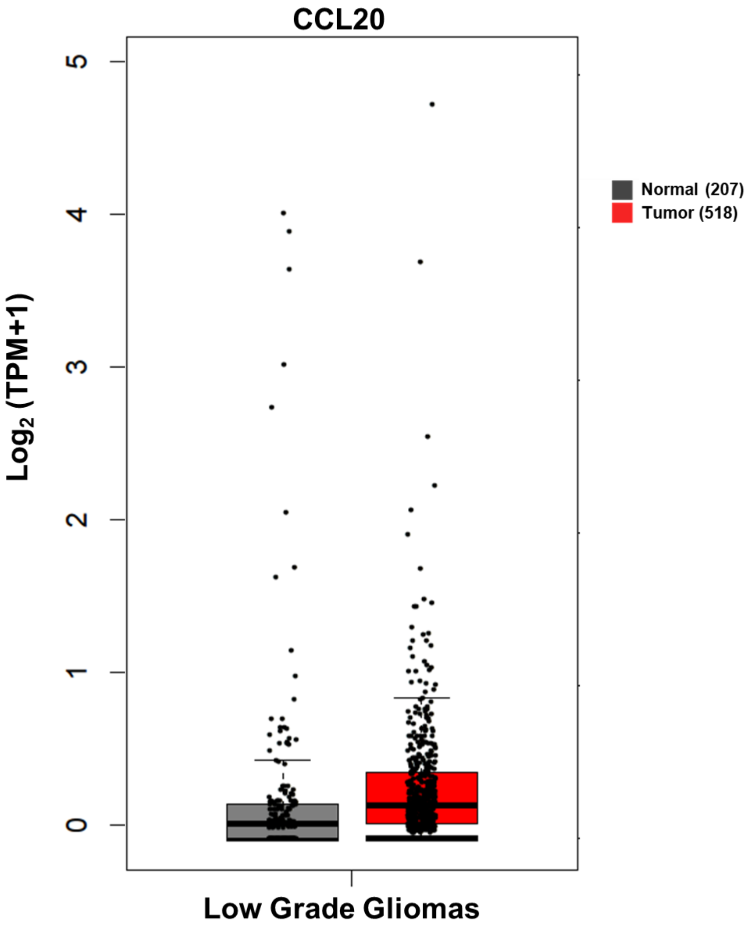


Supplementary Figure 2. CCL20 expression in low grade glioma patients was compared to that of normal brain tissue using GEPIA2. GEPIA2 is an updated version of GEPIA for analyzing the RNA sequencing expression data of 9,736 tumors and 8,587 normal samples from the TCGA and the GTEx projects, using a standard processing pipeline. This tool is developed by Zefang Tang, Tianxiang Chen, Chenwei Li and Boxi Kang of Zhang Lab, Peking University.

### Supplementary Fig. 3

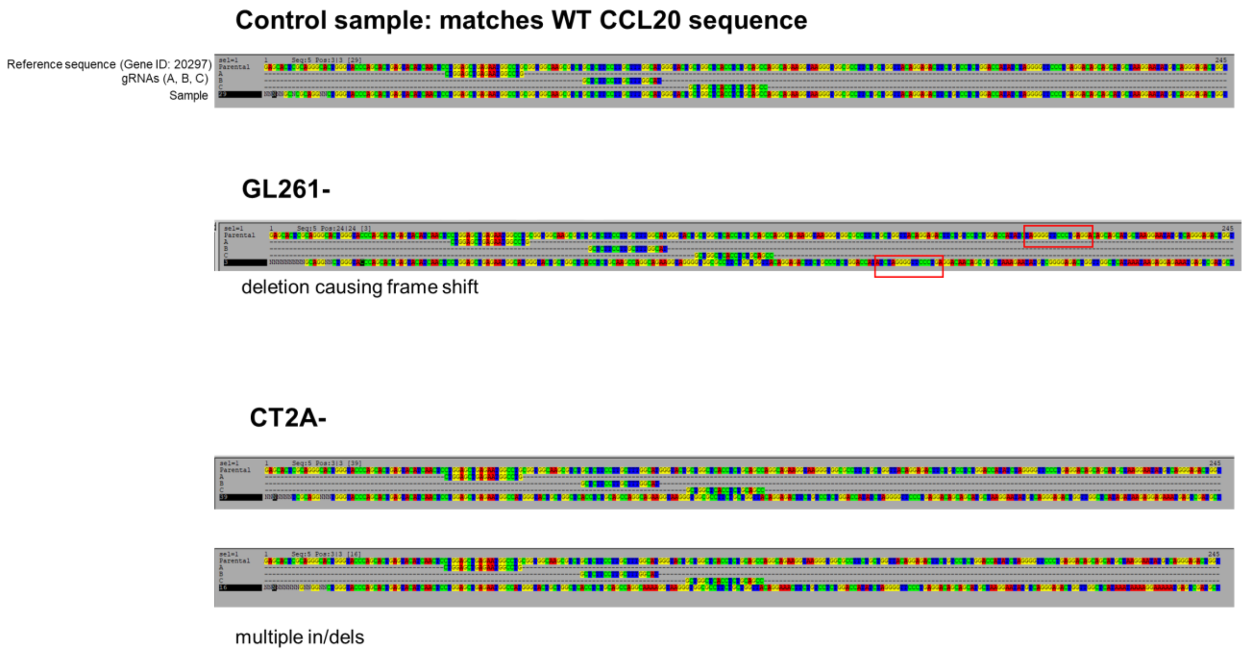


Supplementary Figure 3. Sanger sequencing was performed to verify editing of the CCL20 gene by CRISPR/Cas9 following transfection and clonal expansion (GeneWiz).

### Supplementary Fig. 4

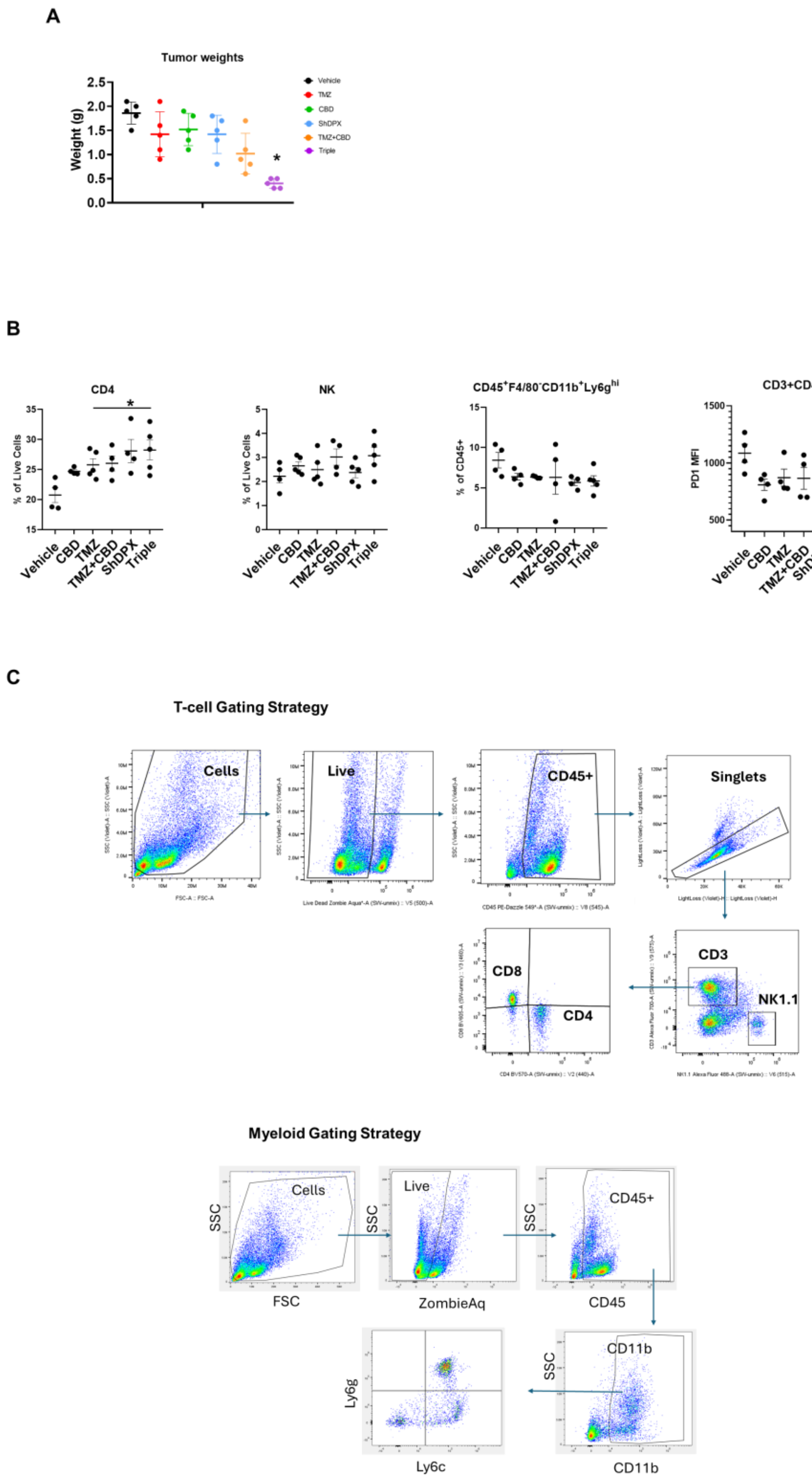


Supplementary Figure 4. Weights of subcutaneous CT-2A tumors were recorded at endpoint (A). Spleens were collected and immunophenotyping of splenocytes for myeloid and T-cell populations was performed (B) using the gating strategy shown (C).

#### Supplementary Fig. 5

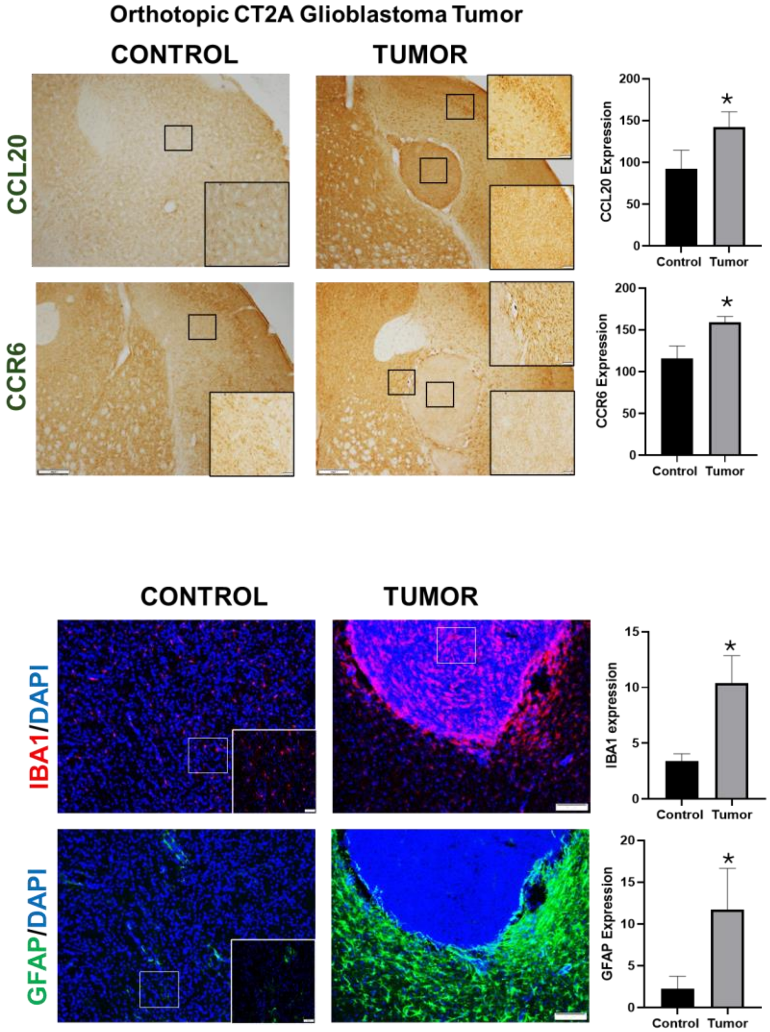


Supplementary Figure 5. Orthotopic CT-2A tumors were collected 14 days post injection. Tumors were cryosectioned and immunostained for CCL20, CCR6, IBA1, and GFAP. Signal was quantified using ImageJ (Student's t test, n=3 *=p≤0.05).

### Supplementary Fig. 6

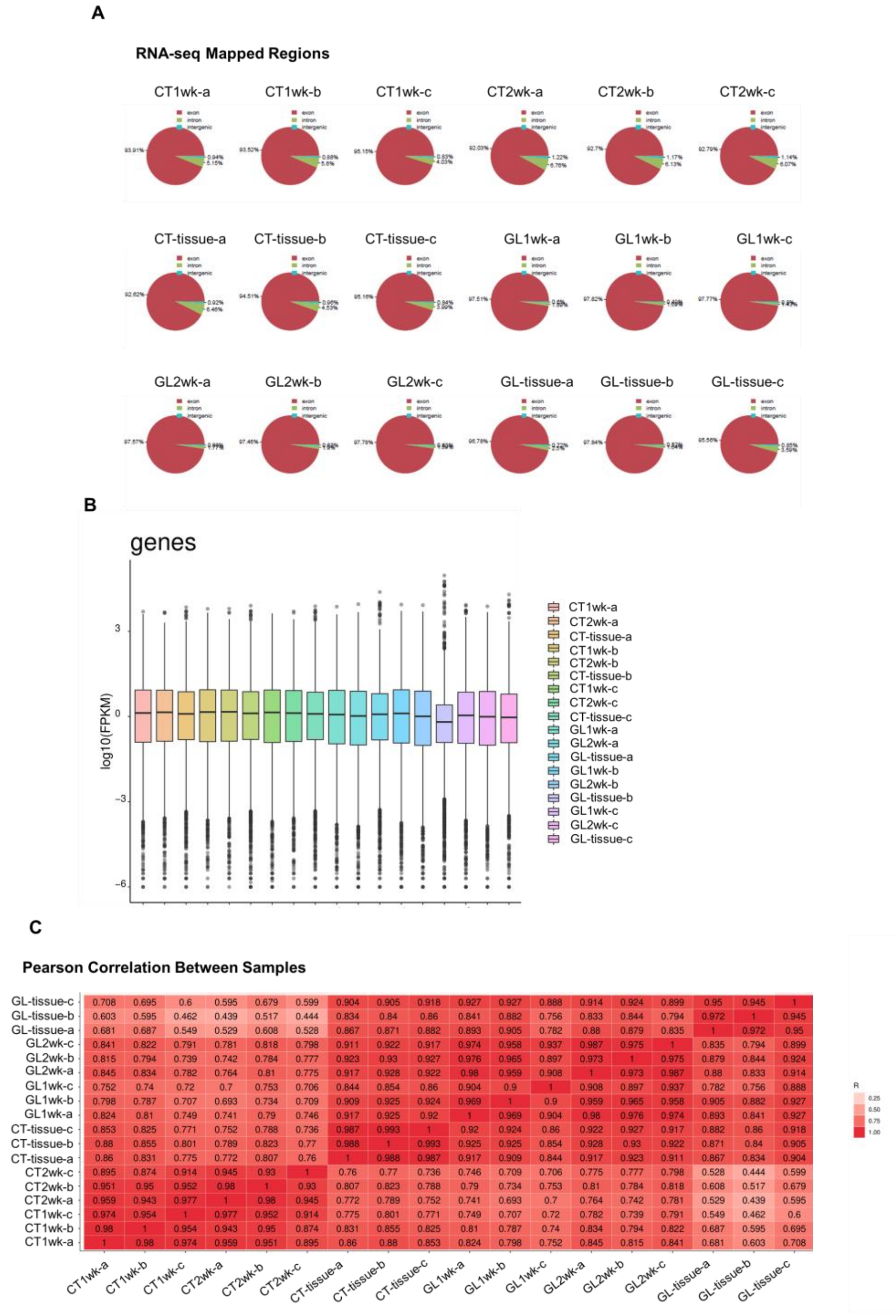


Supplementary Figure 6. Poly-A RNA sequencing of orthotopic CT-2A and GL261 brain tumors and tumor derived organoids was performed by LC Sciences. Proportion of mapped reads (A) and distribution of transcript expression (B) are shown for each sample. Similarity of gene expression between samples was analyzed using a Pearson correlation (C).

#### Supplementary Fig. 7

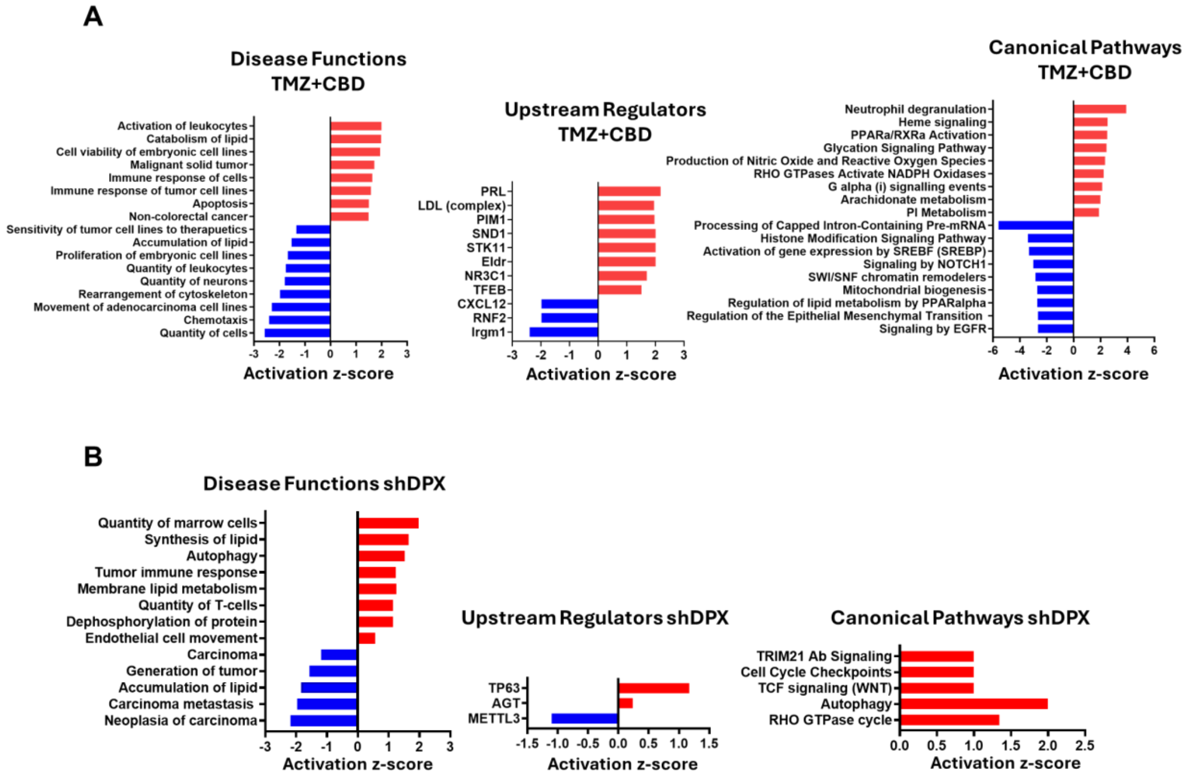


Supplementary Figure 7. IPA pathway analysis of DEPs in CT-2A tumors treated with TMZ+CBD vs vehicle (A) or shDPX vs vehicle (B).

### Supplementary Fig. 8

#### Specialized pro-resolving mediators

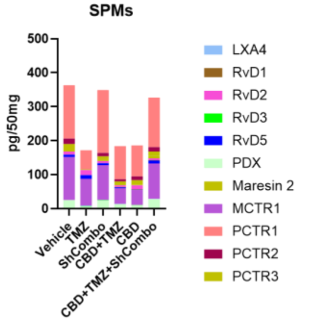


#### Fatty acids

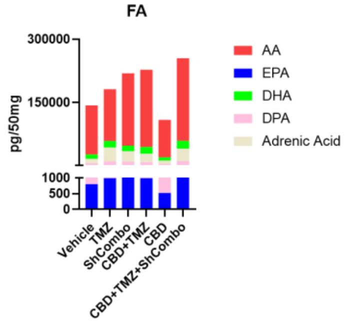


#### Prostaglandins and proinflammatory mediators

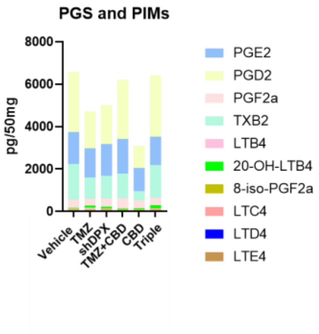


#### proinflammatory mediators

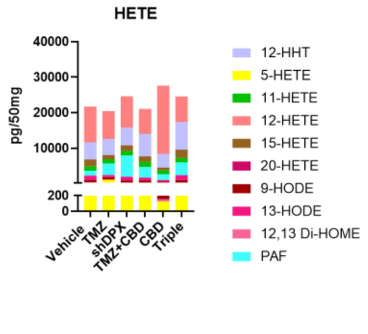


#### Proresolution mediators

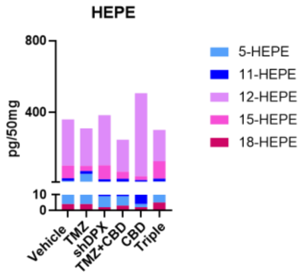


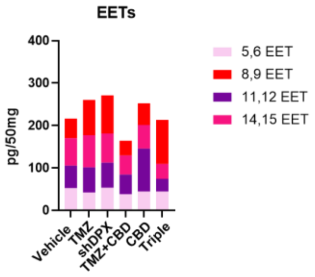


Supplementary Figure 8. Lipidomic analysis of CT-2A tumors.
